# The tumour microenvironment shapes dendritic cell plasticity in a human organotypic melanoma culture

**DOI:** 10.1101/561530

**Authors:** S Di Blasio, M Tazzari, G van Wigcheren, A van Duffelen, I Stefanini, M Bloemendal, M Gorris, A Vasaturo, G Bakdash, SV Hato, J Schalkwijk, IJM de Vries, EH van den Bogaard, CG Figdor

**Author notes:** These authors contributed equally to this work (S Di Blasio, M Tazzari). The Unit of Immunotherapy - Cell Therapy and Biobank, Istituto Scientifico Romagnolo per lo Studio e la Cura dei Tumori (IRST) S.r.l. IRCCS, Meldola (FC), Italy.

## Abstract

The tumour microenvironment (TME) forms a major obstacle in effective cancer treatment and for clinical success of immunotherapy. Conventional co-cultures have shed light into multiple aspects of cancer immunobiology, but they are limited by the lack of physiological complexity. We developed a novel human, organotypic skin melanoma culture (OMC) that allows real-time study of host-malignant cell interactions within a multi-cellular tissue architecture. By co-culturing keratinocytes, fibroblasts and immune cells with melanoma cells, onto a de-cellularized dermis, we generated a reconstructed TME that closely recapitulates tumour growth as observed in human lesions and supports cell survival and function. We demonstrate that the OMC is suitable and outperforms conventional 2D co-cultures for the study of TME-imprinting mechanisms. Within the OMC we observed the tumour-driven conversion of cDC2s into CD14^+^ DCs, characterized by a an immunosuppressive phenotype. The OMC provides a valuable complement to current approaches to study the TME.

## Introduction

Targeting immunomodulatory pathways within the tumour microenvironment (TME) entered centre stage in cancer treatment^1^. Despite promising clinical results of novel cancer immunotherapies, such as immune checkpoint blockade to treat melanoma skin cancer, clinical efficacy is limited and only a minority of patients displays long-lasting clinical responses^2-4^. It is widely accepted that in particular an immunosuppressive TME represents a major hurdle to cancer clearance by immune cells^5,6^. How such an immunosuppressive TME is induced is nowadays intensely studied. Human melanoma models that resemble the complex tissue architectures could improve our understanding of the contribution of the TME during anti-cancer therapy. This will be pivotal for the design of novel strategies that tackle immunosuppressive networks in melanoma to overcome immunotherapy failure^7^.

Over the past decades, a vast array of experimental approaches has been devised to study the melanoma TME, each presenting unique strengths and flaws^8,9^. Those models range from two-dimensional (2D) cultures to whole tissue explants. Albeit informative for basic aspects of cancer biology, 2D culture systems are a poor copy of the *in vivo* cellular environment, as they do not accurately mimic the meshwork of human tissues^10^. Cells in tissues face complex and structurally heterogeneous three-dimensional (3D) architectures and are exposed to a multitude of cellular and extracellular parameters, which influence tumour growth and the ability of stromal and immune cells to orchestrate immune responses locally^7,11^. Tissue explants obtained from a patient’s tumour can be cultured *ex vivo* for microscopic evaluation, or implanted into immunodeficient mice. Tissue explants and patient-derived xenografts (PDXs) retain cell-cell interactions as well as some tissue architecture of the original tumour and are very useful to monitor natural growth of cancer and to investigate tumour heterogeneity^12^. Nevertheless, PDXs lack a functional immune component, limiting their applicability for the study of therapeutic responses to immunotherapy^6^. On the other hand, patient-derived tumour explants can only assess pre-existing tumour immune infiltration (as found at the time of tissue resection), and as such only provide a snapshot. In conclusion, the human melanoma microenvironment including its immune cell components is difficult to mimic using current experimental models.

The potential of a human culture system that accurately mimics the *in vivo* situation, while having the benefit of a laboratory-controlled environment, has inspired researchers to develop 3D models of skin in which skin cancers, such as melanoma, can be propagated (also referred to as skin equivalents or organotypic cultures). In these models, human epidermal cells and melanoma cell lines, with different invasive capacities, are seeded onto fibroblasts-enriched, animal-derived collagen matrices^13-15^. Others developed organotypic cultures based on acellular, de-epidermized human dermis or self-assembled living sheets made with human fibroblasts secreting their own ECM, to obtain a model that more closely resembles the multifaceted skin tissue^16-19^. However, despite important contributions to the field, organotypic skin cultures of melanoma that encompass both stromal- and immunocompetence have never been described^11^.

Intra-tumoural antigen-presenting dendritic cells (DCs) play an important role in stimulating tumour specific cytotoxic T cells, thus driving immune responses against cancer^20,21^. Consensus nomenclature for immune myeloid cells classifies human DCs as conventional DCs (cDCs) and plasmacytoid DC (pDC). cDCs can be further subdivided into cDC1 (CD141^+^ DCs) and cDC2 (CD1c^+^ DCs) subsets^22^. Despite their crucial role in anti-cancer immunity, evidence suggests that DCs in tumours become largely defective and are no longer capable to alert the immune system to cancer. Furthermore, they are often outnumbered by other myeloid cell subsets, such as tumour-associated macrophages (TAMs) and myeloid-derived suppressor cells (MDSCs) that actively suppress the immune system^23-25^. Moreover, we and others have recently described the enrichment of a myeloid cell population in cancer patients that co-expresses markers of monocytes/TAMs (such as CD14, CD163) and cDC2s (CD1c)^26-28^. Given their phenotypic distinction from monocytes and macrophages, these cells are called ‘CD14^+^ DCs’^27^. CD14^+^ DCs are increased in the circulation of advance stage cancer patients and infiltrate both primary and metastatic tumour sites^26-28^. How these CD14^+^ DCs are generated remains yet to be determined.

Building on previous knowledge^19^, in this study we developed an organotypic skin melanoma culture (hereafter referred to as OMC), which contains both stromal and immune components. We validated our OMC by exploring functional plasticity of naturally circulating cDC2s that infiltrate the reconstructed melanoma tissue. Interestingly, we observed that the presence of a tumour within the engineered TME led to an extremely rapid transformation of normal immunostimulatory cDC2s into CD14^+^ DCs, with a phenotype matching their *in vivo* counterpart and an impaired ability to stimulate T cell proliferation. Our results highlight how the newly-generated OMC is instrumental to study tumour-induced events within the TME, that could otherwise not be addressed by static assessments of tissue biopsies.

## Results

### Human organotypic skin melanoma culture mimics natural primary human melanoma lesions

A de-cellularized dermis was used as a scaffold to generate organotypic skin melanoma cultures (OMCs, Fig. 1a). Histochemical evaluation of dermal markers demonstrated that physical decellularization (outlined in the method section) of the human dermal scaffold did not disrupt its complex extracellular matrix architecture (elastin and collagen fibers), and preserved an intact basement membrane (BM) (Suppl Fig 1a,b). Quantitative assessments of the de-cellularized dermis confirmed the lack of cellular (nuclei) and vascular (CD31) components (Suppl Fig 1c-f).

**Figure 1.**
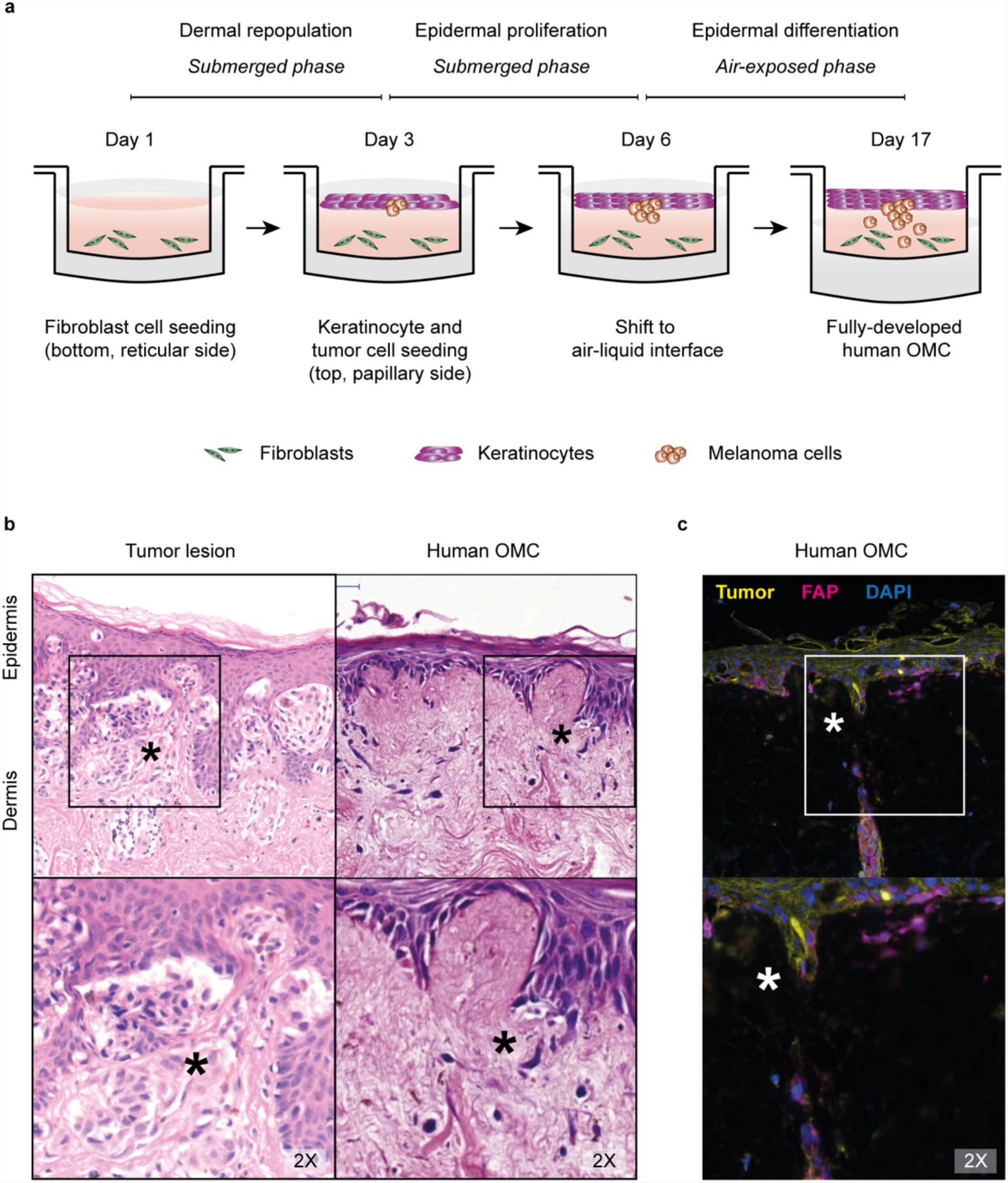
Organotypic human skin melanoma culture generation and characterization. **a,** Overview of the generation of the OMC (prior to immune cell addition). **b,** Histological comparison of tissue sections obtained from primary tumour lesion (left) and OMC (right), showing epidermal and dermal compartments with interspersed tumor nests (asterisks). Hematoxylin-eosin staining, representative pictures. **c,** Multiplex fluorescence immunohistochemistry shows representative area containing fibroblasts (Fibroblast-Associated Protein, FAP^+^ cells, magenta) and melanoma cells Tumor (tyrosinase and SOX10)^+^ cells, yellow). DAPI (blue) indicates nuclei. Original magnification is shown.

To generate a fully developed OMC, primary human fibroblasts (Fbs), keratinocytes (KCs) and melanoma cells are incorporated onto a de-epidermized dermis, and kept under specific culture conditions until a fully differentiated epidermal layer with interspersed tumour nests is formed (Fig 1a). To mimic melanoma growth in the OMC, we co-seeded KCs with melanoma cells on the basal membrane layer, at different KC-to-tumour cell ratios. Immunohistochemistry analysis of model tissue sections and comparison to primary human tumour biopsies showed that the KC-to-tumour cell ratio is critical. At low KC:tumour cell ratios, extensive proliferation of melanoma cells negatively affects the morphology of the epidermal layer. Co-seeding large amounts of fast-diving tumour cells with epidermal cells, caused the tumour cells to outnumber KCs, thereby affecting KC differentiation and preventing the formation of the five characteristic epidermal strata characteristic for fully differentiated skin (Suppl. Fig 2). The optimal cell seeding concentration, defined as the amount of KCs and melanoma cells that preserves epidermal differentiation and morphology, while allowing proliferation of tumour cells to form characteristic compact nests of cells that infiltrate the underneath dermis, was found to be 25 KCs:1 melanoma cell (Fig 1b, Suppl. Fig 2).

**Figure 2.**
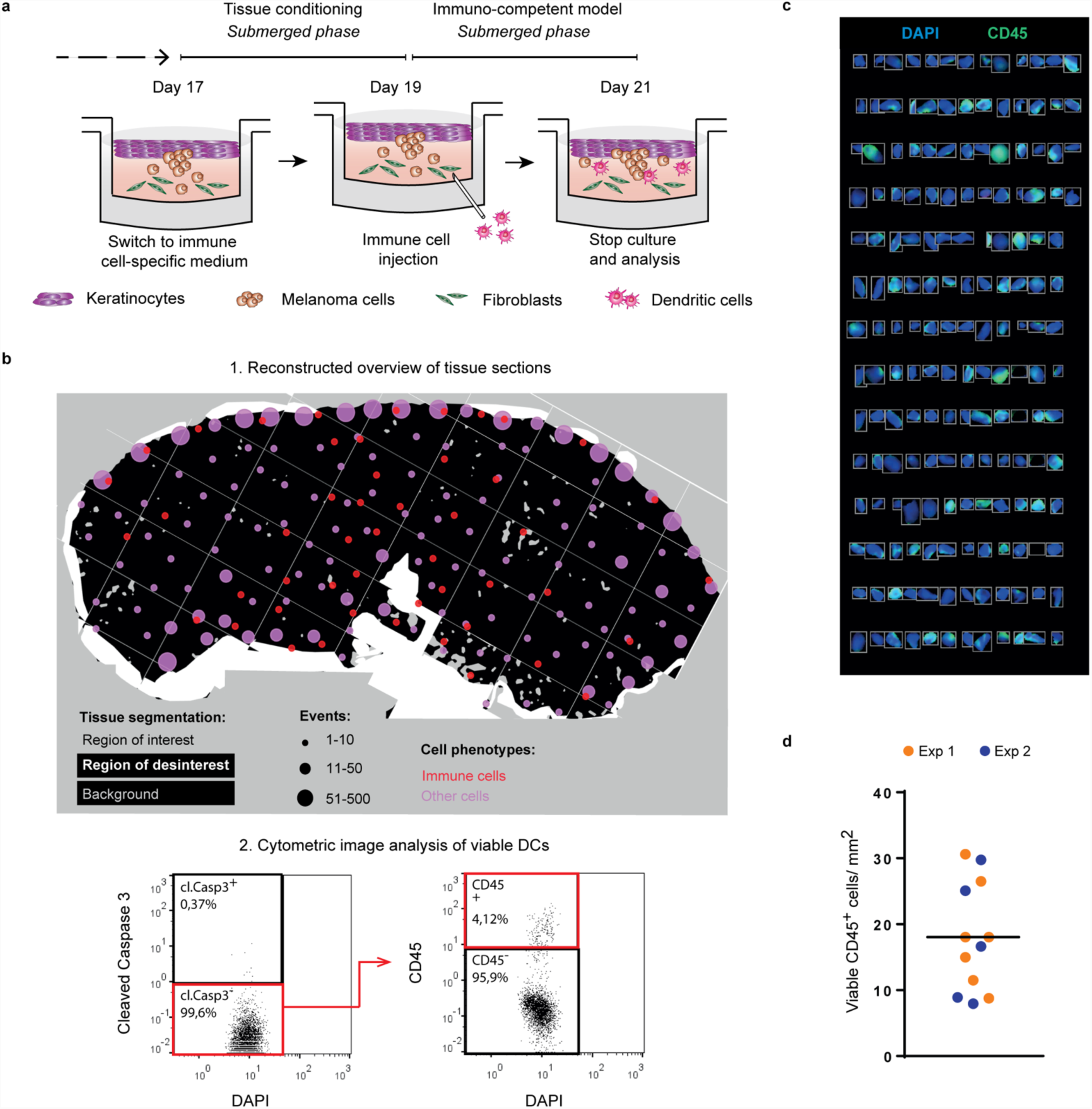
The organotypic skin melanoma microenvironment sustains immune cell survival and distribution. **a,** Overview of the experimental approach used to obtain an immunocompetent OMC. **b,** Tissue classification and DC localization within the selected ROI. Colored dots indicate distribution of immune (red) and other cell types (pink), based on tissue segmentation and positivity score (upper panel, 1); Cytometric image analysis and gating strategy to identify viable DCs (DAPI^+^ Cleaved Caspase 3^-^ CD45^+^ events) (lower panel, 2). **c,** Viable cDC2s as a result of the gating strategy visualized in the original image, showing a representative selection of gated events. DAPI (blue), CD45 (green). **d,** Quantification of viable immune cells/mm^2^ in at least 5 sections per construct, wherein colours indicate independent experiments (n=2).

As shown by multiplex fluorescence immunohistochemistry, melanoma cells and cancer-associated Fbs interact within the OMC (Fig. 1c). Taken together, these observations show that the OMC clearly mimics the invasive growth of tumour cells into the dermis, closely resembling malignancy-associated lesions in human skin.

### Human organotypic skin melanoma microenvironment facilitates tumour-immune cell interactions

A major objective to exploit OMCs is to study human host-tumour interactions, which is notoriously difficult in most xenograft models. In this study, we incorporated a major subset of DCs directly isolated from the peripheral blood (cDC2s, phenotypically defined as CD1c^+^CD14^-^)^26,27^, into the dermal compartment of a fully differentiated OMC (*day 19*, Fig. 2a).

After immunohistochemical staining, the distribution of cDC2s was assessed by reconstructing 20x images into overviews of entire tissue sections (Fig. 2b-d, Suppl. Fig. 3). As a simple metric to quantify the cell density and location within the tissue area (or region of interest, ‘ROI’; Suppl. Fig. 3b), we computed the amount of immune cells and other cells (including stromal and epidermal cells), based on cell segmentation and positivity score using inForm (Fig. 2b, top panel). Enumeration of viable, single cDC2s was obtained with qualitative assessment of signal intensities, analogous to flow cytometry data, for cytometric image analysis as showed in Fig. 2b (bottom panel), followed by microscopic evaluation of gated live, defined as Cl. Cas 3^-^ based on Cleaved-Caspase 3 staining, single immune (CD45^+^) cell (DAPI^+^) events (Fig. 2c). We found that cDC2s were homogeneously distributed throughout the reconstructed melanoma tissue (± 20 cDC2s counted per mm^2^) indicating that immune cells easily migrate into the organotypic culture (Fig. 2d). Altogether, these findings demonstrate that the OMC is a potentially valuable tool to study immune cell behaviour within the TME.

**Figure 3.**
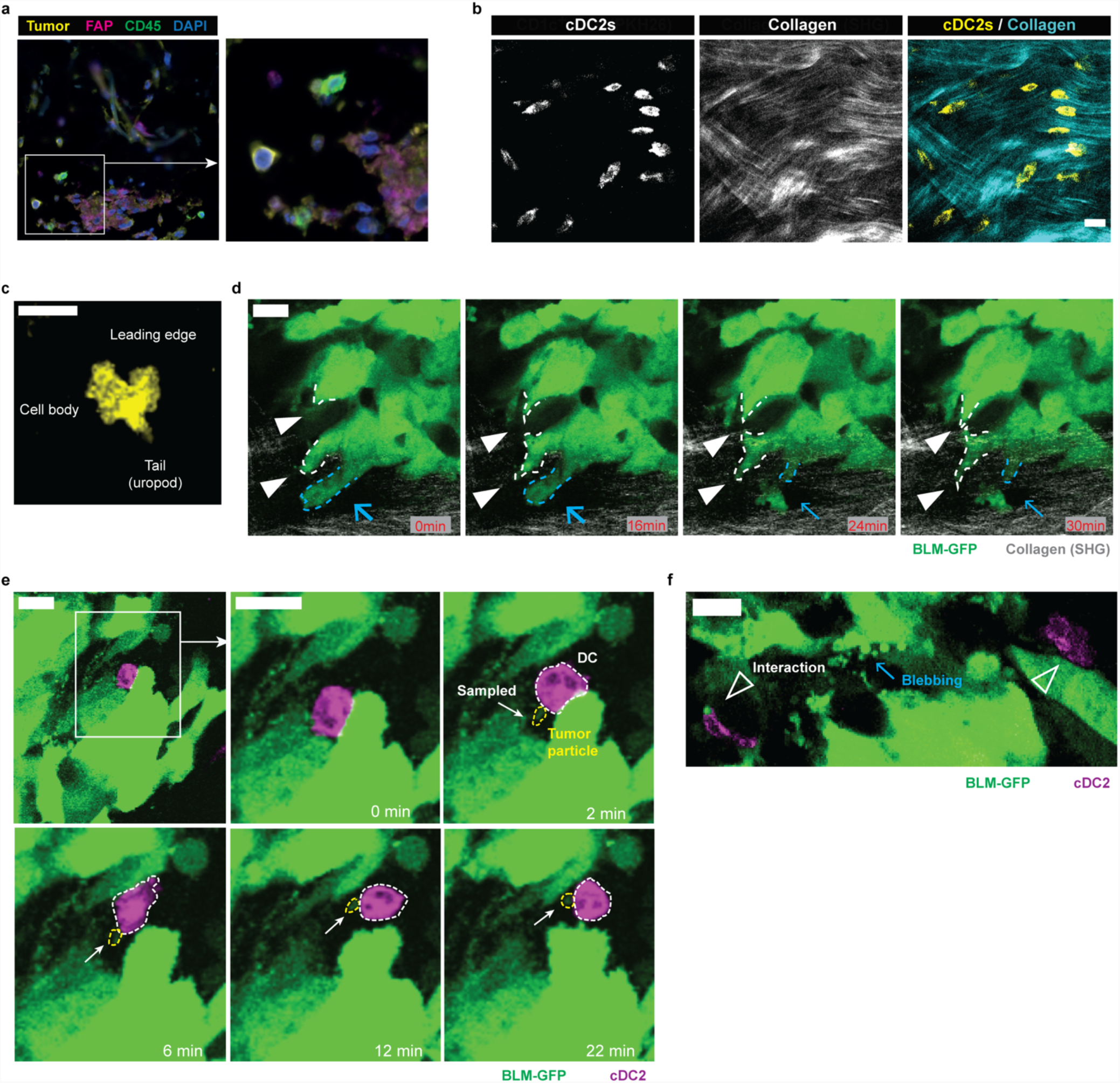
Visualization of cDC2-tumour cell interaction. **a,** Section of OMC showing multiplex fluorescence immunohistochemistry representative area where cDC2s (CD45^+^ cells, green) are found in close proximity to melanoma cells (Tumor (tyrosinase and SOX10)^+^ cells, yellow) and fibroblasts (Fibroblast-Associated Protein, FAP^+^ cells, magenta). DAPI (blue) indicates nuclei. **b,** PKH26 (yellow) and second harmonic generation (SHG, cyan) signals of multiphoton images, acquired with excitation wavelength (λ) of 950nm. PKH26 signal indicates cDC2s; SHG was generated by collagen bundles. **c,** Characteristic amoeboid behaviour of DCs patrolling the environmental niche. **d,** Representative time points during time-lapse recording of melanoma BLM cells expressing GFP [λ (excitation) = 950nm]. BLM cells showed high cellular dynamics (dotted lines; protrusion, *white arrowhead* and retraction, *blue arrow*). **e-f,** Representative time points during time-lapse recording of cDC2s (PKH26)-tumour cell (GFP) interaction [λ (excitation) = 950nm]. (**e**) DC (dotted line, white) sensed and sampled tumour-derived particle (dotted line, yellow). (**f**) Prolonged interaction of DCs with tumour-derived fragments. Tumour cells showed intense membrane dynamics and blebbing. Scale bars, 20μm.

### Dendritic cells interact with melanoma cells and sample tumour-derived particles in a human organotypic skin melanoma microenvironment

Immunohistochemistry end point analysis of fixed OMCs revealed that cDC2s in the dermal compartment were in close proximity with both melanoma cells and cancer-associated Fbs (Fig 3a). To confirm that cDC2s actively interact with cells in their surrounding niche and engage in cell-to-cell interactions with tumour cells, we exploited live two-photon microscopy with a time-lapse setting (Fig 3b-f). Combination of fluorescent signal (live-cell visualization) with second harmonic generation (SHG, elicited by collagen fibre bundles) delivers important information on the 3D anatomy of OMCs. Like in natural skin, collagen fibers were organized in heterogeneous networks, including randomly arranged loose collagen fibers and aligned in more compacted collagen bundles (Fig. 3b, Suppl. Fig. 1a). cDC2s were highly migratory, as evidenced by a typical “hand mirror shape” (Fig. 3c), characteristic of amoeboid motility. cDC2s displayed a leading edge, consisting of multiple dendrites that intercalated between tissue structures, followed by the cell body containing the nucleus, and a posterior tail (uropod). Furthermore, we observed that GFP-expressing melanoma BLM cells (BLM-GFP) were confined by collagen fiber bundles, but showed intense membrane dynamics, evidenced by protrusive and retractile activity (Fig. 3d). Interestingly, cDC2s in our OMC actively interacted with live tumour cells and even sampled tumour-derived cellular microparticles (or blebs), released by the BLM-GFP cells into the ECM (Fig 3e,f and Suppl. Video 1,2). Dynamic DC-tumour cell interactions were observed as early as a few hours after immune cell injection, over periods of at least 20min (Fig 3e). Tumour blebs remained intact as evidenced by the retained cytoplasmic GFP. Taken together, these observations clearly indicate that the OMC represents a promising and valuable platform for accurate investigation of cellular interplay and function, in an *in vivo-*like skin tissue architecture.

### cDC2s convert into CD14^+^ DCs in human organotypic skin melanoma cultures

We and others have previously described the enrichment of CD14^+^ DCs in metastatic melanoma, leukemia and breast cancer patients^26,27,29^. Givein their clinical relevance in different tumor types, we next evaluated whether direct tumor influence in the OMC could convert immunocompetent cDC2s into CD14^+^ DCs. As nowadays cDC2 directly isolated from peripheral blood are routinely used to prepare DC vaccines to treat cancer patients^30^, it would be important to know if these immunostimulatory cells have the potential to become immunosuppressive within the TME.

To test this we studied how the TME of three different melanoma cell lines (BLM, Mel624, A375) modulate the phenotype and function of immunocompetent cDC2s. Tumour-free organotypic skin cultures (OSCs) were generated as controls. To exclude any contamination with other cell types, highly purified cDC2s from healthy donors (purity >98%. Suppl. Fig 4a) were injected into OSCs or OMCs. After two days of culture, organotypic cultures were enzymatically and mechanically digested. Suppl Fig. 4b shows the gating strategy applied for the discrimination of live immune CD45^+^ and non-immune CD45^-^ cells in the digested OSCs and OMCs.

**Figure 4.**
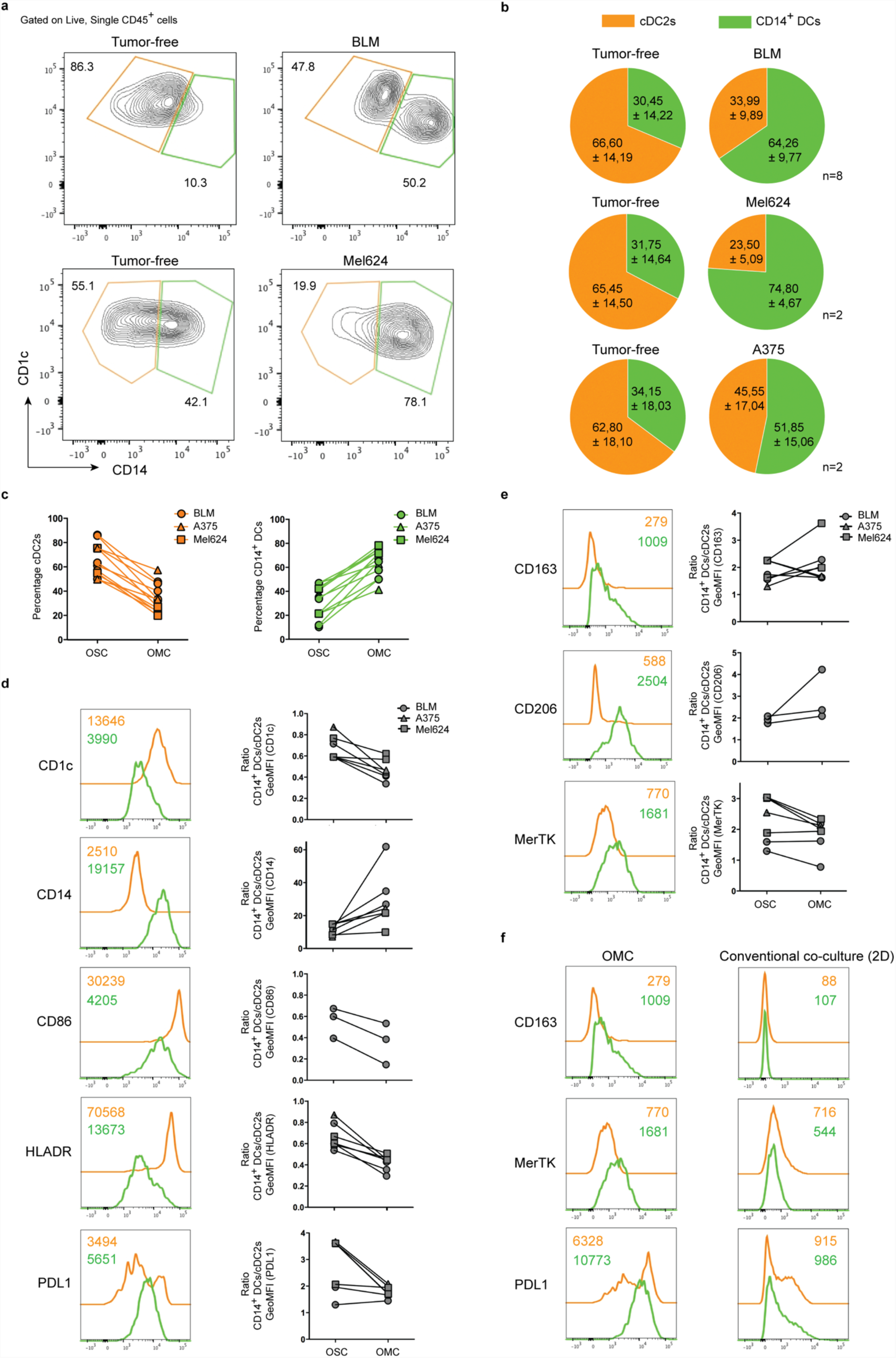
Identification and phenotypic characterization of CD14^+^ DCs isolated from the organotypic human skin melanoma culture. **a,** 2 days after cDC2s injection, organotypic human skin melanoma cultures were digested and cell suspensions were stained. Representative flow cytometry dot plots showing gated cDC2s (CD1c^+^CD14^-^ cells) and CD14^+^ DCs (CD1c^+^CD14^+^ cells) in OSCs (left panels) and OMCs (right panels). Results obtained with two different melanoma cell lines (BLM and Mel624) are shown. Numbers indicate the percentage of gated cells. **b,** Pie charts depicting the frequency of cDC2s and CD14^+^ DCs in OSCs (left) and OMCs (right). Independent experiments were performed with three different melanoma cell lines: BLM (n=8), Mel624 (n=2), A375 (n=2). Data are represented as means ± SD. **c,** Graphs showing the frequency of cDC2s and CD14^+^ DCs in OSCs and OMCs. **d,** Representative overlaid histogram plots showing the expression of the indicated marker in cDC2s and CD14^+^ DCs isolated from OMCs. For each marker a graph reporting the ratio of the GeoMFI between CD14^+^ DCs and cDC2s isolated from OSCs and OMCs is shown. **e,** Representative histogram plots and GeoMFI graphs of macrophage-related markers CD163, CD206 and MerTK in cDC2s and CD14^+^ DCs isolated from OMCs. **f,** Flow cytometry analysis shows the differential phenotype of cDC2s and CD14^+^ DCs isolated from OMCs and conventional tumour-cDC2 co-cultures (2D). Colour legends indicate: cDC2s (orange), CD14^+^DCs (green).

Interestingly, we observed induction of the CD14-monocytic marker on cDC2s isolated from OMCs, compared to cells harvested from OSCs (Fig. 4a). These findings suggest that a monocytic phenotype is induced extremely rapid (within 2 days). Figure 4a shows representative contour plots identifying two distinct populations: cDC2s (CD1c^+^CD14^-^ cells, orange) as originally injected, and cells converted into CD14^+^ DCs (CD1c^+^CD14^+^ cells, green). Pie-charts in Figure 4b illustrate how, in the presence of melanoma, frequencies (mean ± SD) of cDC2s and CD14^+^ DCs change, towards rapid accumulation of the latter and a concomitant prominent reduction of cDC2s in the TME, compared to OSCs. This phenomenon was consistently observed throughout all experiments and independent of the melanoma cell line used (Fig 4c). To further characterize these cells, we assessed whether differences in the expression of the monocytic marker, CD14, reflected additional changes characteristic for CD14^+^ DCs. Tumour-conditioned cDC2s displayed lower CD1c, CD86 and HLADR (GeoMFI CD14^+^ DCs/cDC2s ratio < 1.0) and had higher PDL1 expression (GeoMFI CD14^+^ DCs/cDC2s ratio > 1.0) (Fig. 4d). Interestingly, besides CD14 upregulation, we found that CD14^+^ DCs expressed higher levels of markers typically associated to TAMs: CD163, CD206 and MerTK (GeoMFI CD14^+^ DCs/cDC2s ratio > 1.0) (Fig. 4e). In order to investigate the importance of dimensionality in the tissue microenvironment, we performed intra-donor comparisons (n=4) of cDC2s, isolated from OMCs, to those co-cultured with the same tumour cells in conventional 2D co-cultures. Percentages of CD14^+^ DCs were significantly lower in 2D co-cultures (P=0.0109) compared to those isolated from OMCs (Supplementary Fig 5a,b), providing additional evidence for the importance of 3D organotypic cultures that clearly mimic a skin tissue microenvironment. Moreover, CD14^+^ DCs generated in 2D always failed to up-regulate CD163 and MerTK, and had a much lower expression of PDL1 compared to cells cultured in the OMC (Fig. 4f and Suppl. Fig 5c). Altogether, these data demonstrate that: 1) within the OMC the presence of a tumour rapidly drives cDC2s towards a different myeloid cell subset phenotypically resembling CD14^+^ DCs, 2) the observed phenomenon is tumour-dependent since OSCs lacking tumour cells contains significantly less cells expressing CD14, and 3) 3D multicellular skin cultures are essential, as a conventional 2D co-cultures do not recapitulate such phenotype.

**Figure 5.**
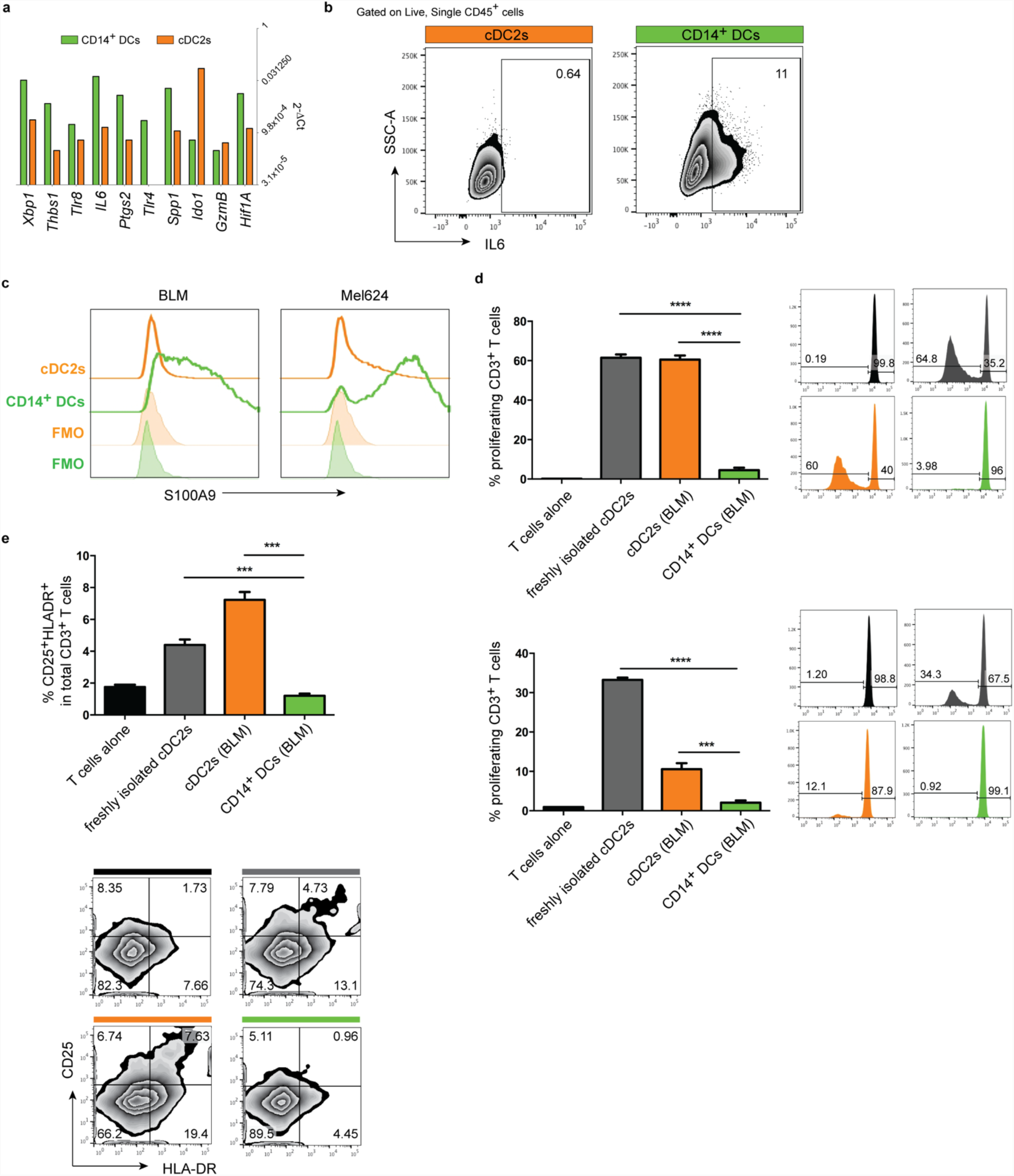
Functional characterization of cDC2s and CD14^+^ DCs isolated from the organotypic human skin melanoma culture. **a,** 2 days after total cDC2s injection, BLM-OMCs were digested and CD1c^+^CD14^-^ (cDC2s) and CD1c^+^CD14^+^ (CD14^+^DCs) subsets FAC-sorted for RNA extraction and molecular characterization by qRT-PCR. Gene expression levels (2^-ΔCt^) for the indicated genes in cDC2s and CD14^+^DCs. ACTB was used as internal reference. **b,** IL-6 production assayed by intracellular cytokine staining in BLM-educated cDC2s, upon stimulation with LPS (1μg/mL) for 6hrs. cDC2s and CD14^+^DCs were identified and gated based on CD1c and CD14 expression in CD45^+^ live single cells. Representative dot plots are shown. Numbers indicate the percentage of gated cells. **c**, Intracellular protein expression of S100A9 in tumour-educated cDC2s and CD14^+^DCs. Representative GeoMFI histograms for BLM and Mel624 are shown. **d,** Proliferation of allogeneic CD3^+^ T cells 5 days after co-culture with FAC-sorted cDC2s and CD14^+^DCs. Right: Representative histogram plots; numbers indicate the percentage of gated cells. Left: Bar graphs (n of replicates = 3, means ± SEM). Two representative experiments out of three are shown. **e,** Autologous T cells were co-cultured with cDC2s and CD14^+^DCs for 5 days and then stained for HLADR, CD25, CD3, CD8 and live/dead marker. Percentage of activated CD25^+^HLDR^+^ cells in total CD3^+^ T cells is reported. Bar graphs (n of replicates = 3, means ± SEM). One representative experiment out of three is shown. Colour legends indicate: cDC2s (orange), CD14^+^DCs (green).

### Melanoma-induced CD14^+^ DCs display immunosuppressive function

To functionally characterize tumour-induced CD14^+^ DCs, we next performed qRT-PCR on cDC2s and CD14^+^ DCs FAC-sorted 18 hours post-injection. We confirmed at the transcriptomic level that CD14^+^ DCs from OMCs express CD14, CD163 and CD206 genes (data not shown), consistent with protein surface expression levels as measured by flow cytometry (Fig. 4d). Interestingly, we observed that these CD14^+^ DCs generated within the OMC also expressed higher levels of markers previously linked to human myeloid cells endowed with suppressive activity: *XBP1, THBS1, IL6, PTGS2 and HIF1A* (Fig. 5a, Suppl. Fig 6a)^31,32^. At the protein expression level, we confirmed that only CD14^+^ DCs promptly produced IL-6 (0.64% vs 11%), upon TLR4 stimulation (Fig. 5b). Intriguingly, we observed that these cells could also be distinguished based on their S100A9 expression (Fig. 5c), underlining another similarity to already described regulatory myeloid cell features^33^. Of note, the immunosuppressive molecule IDO1, often found in association with tolerogenic DCs^34^ was predominantly expressed in cDC2s that still retained a clear DC phenotype upon interaction with tumour cells (Fig. 5a). This suggests that within OMCs, injected cDC2s follow two different fates: 1) they become tolerogenic cDC2s; 2) they convert to CD14^+^ DCs. To further investigate if these cDC2s also functionally reverted to immunosuppressive myeloid cells, we performed mixed lymphocyte reactions to test their immunostimulatory potential. cDC2s that acquired CD14 during culture, were far less capable of stimulating allogenic CD3^+^ T cells when compared to cDC2s that lacked CD14 (Fig 5d). Similarly, they were significantly less able to induce autologous T cell activation (Fig. 5e). Of note, we observed that CD14^+^ DCs, isolated from OSCs, were already less stimulatory than cDC2s, but the presence of the tumour further decreased their ability to stimulate T cell proliferation and activation (Suppl. Fig. 6b,c). Collectively, our data show that cDC2s that acquire CD14 within OMCs, represent a functional distinct myeloid population induced by the TME.

**Figure 6.**
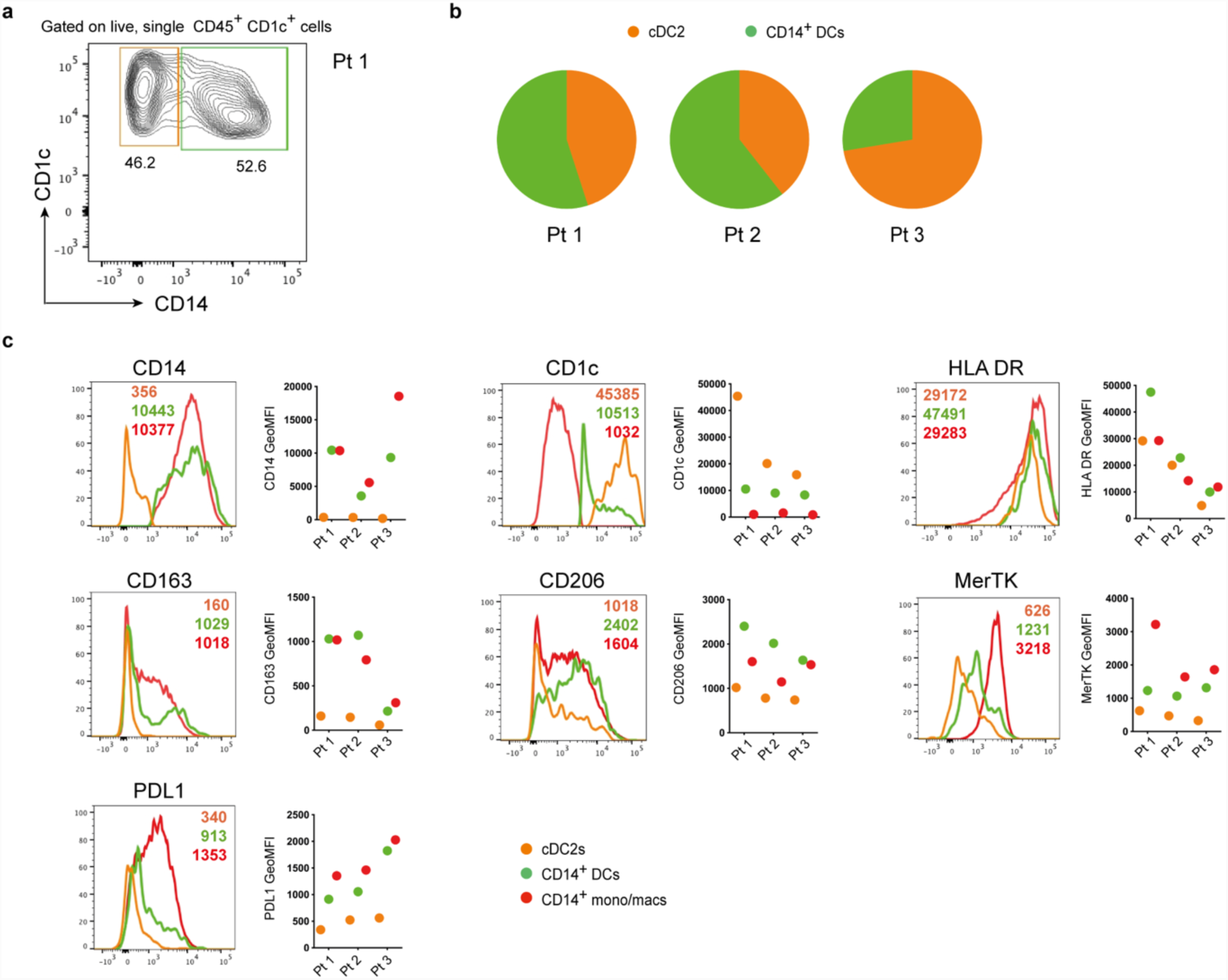
Human melanoma lesions contain CD14^+^DCs phenotypically close to those recapitulated in the organotypic human skin melanoma culture. **a,** Human melanoma lesions were digested and stained (n=3). CD1c^+^CD14^-^ (cDC2s), CD1c^+^CD14^+^ (CD14^+^DCs) were identified and gated based on CD1c and CD14 expression in CD45^+^CD1c^+^ live single cells. A representative dot plot from one patient (Pt) is shown. **b,** Pie charts report percentages of cDC2s and CD14^+^DCs across different melanoma patients (n=3). **c,** Graphs showing GeoMFI’s of the phenotypic analysis of cDC2s, CD14^+^DCs and CD14^+^ mono/macs defined within intra-tumor CD45^+^CD11c^+^ live single cells from melanoma patient samples. Representative histogram plots for each indicated marker are shown. Colour legends indicate: cDC2s (orange), CD14^+^DCs (green), CD14^+^ mono/macs (red).

### CD14^+^ DCs generated in the organotypic human skin melanoma model resemble those found in melanoma lesions

Finally, we set out to compare CD14^+^ DCs induced in the OMC with myeloid cells, as isolated from melanoma lesions of stage IV metastatic melanoma patients. Using flow cytometry analysis, we identified CD14^+^ DCs (green) and cDC2s (orange), similarly to what previously observed within OMCs (Fig 6a). Pie charts in panel b summarize the results obtained across three different patients. Further phenotypic analyses revealed the macrophage-like nature of this *ex vivo* CD14^+^ DCs characterized by a higher expression of CD163, CD206 and MerTK (Fig. 6c). Histograms in panel c show that *ex vivo* CD14^+^ DCs, identified within intra-tumoral CD45^+^CD11c^+^ cells (Suppl Fig. 7), have a phenotype that is closer to that of CD14^+^ mono/macs (red), rather than cDC2s. Moreover, we could confirm that also *ex vivo* CD14^+^ DCs expressed higher levels of PDL1 when compared to cDC2s. Overall, this *ex vivo* analysis of tumour biopsies highlights the *in vivo* similarity of CD14^+^DCs with those reprogrammed within OMCs.

## Discussion

Recently, tumour organoids have received much attention^35,36^. Organoids have the advantage of retaining both histological and mutational features of the original tumour, and can be propagated for extended periods of time, thus facilitating extensive experimenting. Yet, they fail to represent the multicellular composition of the TME^37^. In particular, the inclusion of stroma and tumour-infiltrating lymphocytes has been only recently described^37^. So far organoid cultures have mainly been generated from epithelial cancer types^35,36,38^, leaving the more complex skin tissue, where primary melanoma resides, still uncovered. In the context of human melanoma, local tumour growth was mimicked in *in vitro* skin equivalents, which however lacked an immune compartment^16^. With the aim of filling this gap, we generated a novel human multicellular skin microenvironment, amenable to controlled experimental manipulation, suitable to explore cancer-driven immune cell modulation within a complex tissue architecture. Similar to organoids, the OMC we developed, reproduces the spatial distribution of cells that naturally constitute the skin. In this respect, we observed that an optimal KC:tumour cell ratio is essential to ensure the formation of the epidermal layer, while allowing proliferation of tumour cells into nests that infiltrate the underneath dermis.

Organoids and tumor spheroids are developed using synthetic and animal-derived scaffolds. However, these matrices, while ensuring the required tissue stiffness, may otherwise act as a source of foreign antigens^39^. Thus, de-cellularized human tissue like the one here employed, or synthetic matrices^40^ potentially represents better alternatives for establishing an immunocompetent TME. Interestingly, besides the illustrated use of primary naturally circulating DCs, we assessed the feasibility of culturing T lymphocytes within the OMC, and found that the percentage of live CD3^+^CD4^+^ and CD3^+^CD8^+^ cells was not affected by the culture conditions (Suppl. Fig. 8). Remarkably, the rapid changes (2 days) we observed on cDC2s cultured in OMCs, did not require the addition of exougenous growth factors or cytokines, which could *per se* potentially alter cell phenotype and function. Nevertheless, whether the use of supplemented culture media is required for longer culture periods or different immune cell subsets, remains to be explored.

As previously reported for patient-derived organoids and tumor spheroids, we also explored the use of the de-cellularized dermis as a scaffold for infusing material directly isolated from melanoma biopsies (Suppl. Fig. 9). The analysis of the lymphoid and myeloid populations, by means of multiplex IHC, assessed vitality and cellular distribution. Further experiments are desirable to confirm suitability of the human de-cellularized dermal scaffold to maintain tumour and immune representativeness of the original lesion.

The intra-tumoural infiltration and activation of myeloid cells, such as DCs, has clinical relevance across different tumour types^21^. Thus, understanding the mechanisms that regulate the fate of individual DC subsets within the TME will be pivotal to define strategies that revert DC immunosuppression while simultaneously enhancing their activation^41,42^. Importantly, the behaviour of antigen-presenting cells in a tumor organoid-like context has never been addressed.

We validated our model by following the response of circulating cDC2s within the reconstructed TME. cDC2s are phenotypically defined as CD1c^+^CD14^-^ and characterized by the ability to stimulate cytotoxic T cell responses, exemplified by their use in DC vaccination protocols^30^. By combining imaging, flow cytometry and gene expression profiling we documented the interaction of cDC2s with stromal and melanoma cells within the OMC, and attested the local conversion of cDC2s into a distinct myeloid cell population. Of note, this tumor-induced myeloid subset, characterized by the decreased expression of the DC marker CD1c, and the concomitant acquisition of monocyte/TAM markers (CD14, CD163, CD206 and MerTk), is consistent with CD14^+^ DCs already described in literature in the context of cancer-related inflammation^27,28^. These findings argue in favour of the ability of the OMC to recapitulate *in vivo*-like phenomena. While the current literature indicates that CD14^+^ DCs arise from monocytes^22^, we here suggest a new route of melanoma-mediated conversion of cDC2s into CD14^+^ DCs. Such cDC2s-inherent phenotypic plasticity is not counterintuitive, if we reason that single cell-RNA sequencing recently revealed the unappreciated heterogeneity of blood cDC2s, isolated from healthy individuals. Indeed, a cell cluster characterized by a unique inflammatory gene signature, close to that of CD14^+^ monocytes, has been identified within the CD14-negative cDC2 subset^43^. Functionally, the melanoma-induced CD14^+^ DCs described here, express genes (such as *SSP1*, *PTGS2*, *IL6*) previously associated with immunosuppressive myeloid cells^31,32^ and, like monocytes and macrophages, have poor T cell stimulatory ability. The evolution of conventional DCs into regulatory macrophage-like cells has been proposed in murine models^44,45^, but to our knowledge this is the first study that reports the ability of human, naturally-occurring, mature cDC2s to be reprogrammed into CD14^+^ DCs. Indeed, human *in vitro* studies investigating this phenomenon were performed using monocyte-derived DCs (moDCs), which represent a poor surrogate of *in vivo* DCs^46,47^. It will now be critical to unravel the molecular drivers of the immunostimulatory cDC2s transition into regulatory CD14^+^ DCs, within tumour tissues. Of note, albeit the three melanoma cell lines, employed for the generation of the organotypic skin model, present differences in gene expression and secretome profiles (data not shown), we so far did not observe dissimilarities in their ability to promote cDC2 conversion.

Our results also support the hypothesis that the multicellular dimensionality of the TME plays a critical role in shaping cell phenotypes^7,11^. We here showed that the phenotype of tumour-induced CD14^+^ DCs was not obtained in (parallel intra-donor) 2D co-culture experiments. In particular, although some expression of CD14 was observed, these cells failed to express the TAM markers, CD163 and MerTK.

The *in vitro* differentiation of myeloid cells to acquire TAM traits usually requires ≥3 days^48,49^. In this respect, we believe that the rapid cDC2s conversion (2 days) could be explained by the fact that tumour and immune cells are not added in culture simultaneously, as usually performed in classical 2D co-culture experiments, but instead immune cells are exposed to an already tumour-conditioned skin microenvironment. This certainly leads to a stronger tumour-mediated effect and more closely resembles the natural cDC2s entry into tumour tissues *in vivo*.

Intriguingly, CD14^+^ DCs developed within the organotypic melanoma cultures shared important phenotypic similarities with CD14^+^ DCs infiltrating human melanoma lesions, and express higher levels of TAM-related markers (CD206, MerTK and CD163), compared to blood circulating CD14^+^ DCs, we previously reported to be enriched in melanoma patients showing a lower immunological response to DC-based vaccines^26,29^. Those results demonstrate the added value of the OMC, with respect to classical 2D co-culture cell systems. Our data seem suggest that the de-cellularized human-derived dermis repopulated with stromal cells, not only provides the correct extracellular space and guidance for cells to migrate, but also offers the mechanical tissue stiffness as observed *in vivo*, and which likely contributes to TAM differentiation^50^.

*Ex vivo* tumour-infiltrating CD14^+^ DCs are characterized by the expression of the T cell inhibitory molecule PDL1. Interestingly, the up-regulation of this key immune escape molecule was higher in tumour-educated myeloid cells within the OMC compared to those developed in classical monolayer co-cultures. In this respect, it would be of interest to use such a model to assess the dynamic modulation of immune checkpoint inhibition.

We are aware that, like most organotypic culture systems, our OMC is limited by the absence of blood vessels and lymphatics, and thus events such as angiogenesis and leukocyte extravasation cannot be mimicked. In order to overcome this caveat, attempts have been made to engineer collagen matrices containing a microvasculature network of endothelial cells, or to integrate microfluidics with tissue engineering to mimic the function of native microvessels^51,52^. Those approaches are still in their early infancy and further development is needed^53,54^. However, we believe that our model could be employed to assess T cell infiltration, for the investigation of mechanisms governing inflamed *versus* not inflamed tumour tissues.

In the present study we describe the generation, characterization and one potential application of a novel OMC. In contrast to patient-derived tumour spheroids, this system does not aim to assess pre-existing immune infiltration^55^, but should be viewed as a fully titratable *in vivo*-like environment for the advanced investigation of tumour-immunological mechanisms. Overall, we believe that the proposed OMC will contribute to the identification of candidate genes and molecules involved in immune escape processes. Ultimately, this might lead to the design of novel therapeutic targeted approaches addressing the melanoma microenvironment.

## Materials & Methods

### Cell culture, transduction and stable cell line development

Human melanoma cells (BLM, BLM-GFP, Mel624 and A375) were tested to be mycoplasma-free, authenticated by ATCC and maintained in Dulbecco’s modified Eagle’s medium (DMEM, Gibco), supplemented with 5% fetal calf serum and 5% CO2 humidified air at 37°C. The Lenti6/Block-iT-shScramble (GFP) vector was a kind gift of Prof. Peter Friedl (RIMLS, The Netherlands). The sequence of this construct does not match any known mammalian genes. BLM cells were infected with lentiviral vector and (10µg/ml) polybrene and incubated at 37°C, 5% CO2, overnight. Then medium was refreshed and cells were analyzed after 72h of treatment. A stable cell line was selected with 5µg/ml blasticidin.

### Isolation of human blood immune cells

Peripheral blood mononuclear cells (PBMCs) were isolated from buffy coats obtained from healthy volunteers (Sanquin) and purified via centrifugation over a Ficoll density gradient (Axis-Shield) in SepMate tubes (Stemcell technologies). cDC2s (CD1c^+^) cells were purified with magnetic cell sorting (MACS) from healthy donor PBMCs using the CD1c (BDCA1) DC isolation kit (Miltenyi Biotec). To obtain a CD1c^+^CD14^-^ population, a pre-depletion step of monocytes using CD14-MACS microbeads (Miltenyi Biotec) was included in the original manufacturer’s protocol. DC purity was assessed by staining with primary directly-labelled antibodies: anti-CD1c, anti-CD14 and anti-CD20 Abs (Supplementary Table 1). Purity levels higher than 98% were achieved, determined by flow cytometry (Suppl. Fig. 4a). For *live* imaging experiments, cDC2s were pre-labelled with the membrane dye PKH26 (Sigma-Aldrich) according to manufacturers’ protocols and resuspended in X-VIVO-15 medium (Lonza) supplemented with 2% human serum (HS, Sanquin). Autologous or allogenic CD3^+^ T cells used in T cell co-culture experiments were isolated using the Pan T cell isolation kit (Miltenyi Biotec) following manufacturer’s instruction.

### Isolation of cellular and extracellular matrix human skin components

Generation of de-epidermized, de-cellularized dermis and isolation of human primary keratinocytes from human abdominal skin, derived from donors who underwent surgery for abdominal wall correction, was performed as previously described^56^. Briefly, human skin was incubated for five to ten minutes in phosphate-buffered saline (PBS) at 56°C to allow separation of the epidermis from the dermis. A de-epidermized human dermis was obtained by incubating the dermis for one month in PBS containing gentamicin (0.5 mg/ml; Life Technologies, Inc.) and antibiotic/antimycotic (Life Technologies, Inc.) at 37°C. Continuous cycles of freezing and thawing ensured the depletion of all living cells in the dermis. Punches were prepared from this de-epidermized dermis using an 8-mm biopter, exposed to additional freezing/thawing cycles and frozen for further use. Keratinocytes were isolated from the epidermal layer by trypsin treatment for 16 to 20 hours at 4°C, and then cultured in the presence of irradiated (3295 cGy for 4.10 minutes) mouse fibroblasts 3T3 cells. 3T3 cells were seeded at a concentration of 3×10^4^ cells per cm^2^ in Greens medium, which consisted of two parts Dulbecco’s modified Eagle’s medium (Life Technologies), one part of Ham’s F12 medium (Life Technologies), 10% fetal bovine serum (Hyclone), L-glutamine (4 mmol/L; Life Technologies, Inc.), penicillin/streptomycin (50 IU/ml; Life Technologies, Inc.), adenine (24.3 μg/ml; Calbiochem, San Diego, CA), insulin (5 μg/ml; Sigma, St. Louis, MO), hydrocortisone (0.4 μg/ml; Merck, Darmstadt, Germany), triiodothyronine (1.36 ng/ml, Sigma), cholera toxin (10^-10^ mol/L, Sigma). The next day keratinocytes were added at a concentration of 5×10^4^ cells per cm^2^. Keratinocytes-3T3 cells co-cultures were maintained for three days in Greens medium. Thereafter, medium was replaced by Greens medium containing epidermal growth factor (EGF, 10 ng/ml; Sigma), cells were expanded until 90% confluence and stored in the liquid nitrogen. Keratinocytes were used at passage one or two in the model. Adult human dermal fibroblasts were purchased by ATCC and maintained in Fibroblast medium (3:1 DMEM: Ham’s F12 Nutrient Mixture, Gibco) supplemented with 10% FCS, and expanded until 80% confluence. Passages three to nine were used for the experiments.

### Generation of the human organotypic skin melanoma culture

To generate the human organotypic skin melanoma culture (OMC), 8mm punch biopsy of de-epidermized, de-cellularized dermis was placed carefully basal membrane side down on a transwell insert in a 24-well plate (24-wells ThinCert, Greiner Bio-One) and 0.25×10^6^ fibroblasts were seeded onto the reticular dermal side (opposite to papillary side and basal membrane) of a de-epidermized dermis, via centrifugal force (day 1), in Fibroblast medium (3:1 DMEM: Ham’s F12 Nutrient Mixture, Gibco) supplemented with 10% FCS. The plate was centrifuged for one hour at 500rpm. After centrifugation, the de-epidermized dermis was maintained in Fibroblast medium for two days at 37°C, to allow cellular proliferation and distribution through the structural collagen bundles and elastin fibers. After two days of culture, the repopulated de-epidermized dermis was turned basal membrane side up in the transwell insert and keratinocytes were seeded together with melanoma cells (BLM, Mel624 or A375) onto the papillary side of the dermal scaffold (day 3). The OMC was cultured submerged for three days in PONEC medium containing 5% serum (PONEC 5% medium), to allow proliferation of keratinocytes and tumour cells (day 6). PONEC 5% medium consists of two parts Dulbecco’s modified Eagle’s medium (Life Technologies, Inc.), one part of Ham’s F12 medium (Life Technologies, Inc.), 5% calf serum (Hyclone), L-glutamine (4 mmol/L; Life Technologies, Inc.), penicillin/streptomycin (50 IU/ml; Life Technologies, Inc.), adenine (24.3 µg/ml; Calbiochem, San Diego, CA), insulin (0.2 µmol/l; Sigma, St. Louis, MO), hydrocortisone (1 µmol/l; Merck, Darmstadt, Germany), triiodothyronine (1.36 ng/ml, Sigma), cholera toxin (10^-10^ mol/L, Sigma), ascorbic acid (50µg/ml; Sigma). Thereafter, the OMC was shifted to the air-liquid interface and cultured for eleven days in PONEC medium without serum, supplemented with keratinocyte and epidermal growth factors (PONEC 0% medium), during which a fully differentiated epidermal layer is formed (day 17). PONEC 0% medium consists of two parts Dulbecco’s modified Eagle’s medium (Life Technologies, Inc.), one part of Ham’s F12 medium (Life Technologies, Inc.), L-glutamine (4 mmol/L; Life Technologies, Inc.), penicillin/streptomycin (50 IU/ml; Life Technologies, Inc.), adenine (24.3 µg/ml; Calbiochem, San Diego, CA), L-serine (1mg/ml; Sigma), L-carnitine (2 µg/ml, sigma), bovine serum albumin lipid mix (palmitic acid 25 µmol/l; arachidonic acid 7 µmol/l; linoleic acid 15µmol/l; vitamin E 0.4µg/ml; all from Sigma), insulin (0.1 µmol/l; Sigma, St. Louis, MO), hydrocortisone (1 µmol/l; Merck, Darmstadt, Germany), triiodothyronine (1.36 ng/ml, Sigma), cholera toxin (10^-10^ mol/L, Sigma), ascorbic acid (50 µg/ml; Sigma), keratinocyte growth factor (5 ng/ml, Sigma), epidermal growth factor (2 ng/ml, Sigma). At this point, the OMC was conditioned for two days with X-VIVO-15 medium (Lonza) supplemented with 5% human serum (Sanquin), to sustain survival of immune cells in the reconstructed microenvironment (day 19). *Ex-vivo* culture of immune cells, which can be freshly isolated from peripheral blood circulation or separated from the tissue they infiltrate, can be a critical procedure due to the fragile nature of those cell types when dissected from their original microenvironment. At day 19, 0.1 to 0.5 x10^6^ cDC2s were microinjected into the dermis using a microneedle array system (kindly provided by Nanopass Technology Ltd, Israel). The microneedle array holder was connected to a manual syringe pump (1ml, BD). Microinjection was performed using air pressure. Care was taken not to allow the microneedles to rupture the regenerated skin. Immunocompetent OMCs were cultured for additional two days in X-VIVO-15 medium plus 5% human serum. For experiments that assessed DC phenotype and function, melanoma cells were incorporated 1) together with Fbs onto the dermal scaffold (day 1), and 2) co-seeded with keratinocytes onto the basal membrane (day 3). The double seeding procedure of melanoma cells was performed to guarantee efficient melanoma conditioning of the reconstructed microenvironment, in a way that would more closely recapitulate *in vivo* tumour-associated tissues.

### Patient material

Tumour specimens were collected from stage IV metastatic melanoma patients. The following cases were analyzed: Pt 1, metastasis at the right upper leg; Pt 2, metastasis at the right adrenal gland, Pt 3 liver metastasis. The study was approved by our Institutional Review Board and written informed consent was obtained from all patients.

### Tumour dissociation

Melanoma cell suspensions were obtained from tumour sample of patients who underwent surgery by enzymatic and mechanic digestion using the gentleMACS Dissociator (Miltenyi, Bergisch-Gladbach, Germany). Briefly, tumour specimens were minced under sterile conditions into small pieces and digested over 1 h at 37 1C following the gentle MACS Dissociator protocol (Miltenyi). The resulting cell suspension was filtered through a 70-μm mesh (BD Biosciences, San Jose, CA, USA), the red blood cells were lysed, and the cell suspension was washed with RPMI. Cells were stored in liquid nitrogen until use. The same approach was used to process OMCs for the isolation and analysis of immune cells.

### Immunohistochemistry and manual digital cell density analysis

Slides of 4 or 6-µm thickness were cut from formalin-fixed, paraffin-embedded (FFPE) primary melanoma tissue blocks and OMCs. Hematoxylin-Eosin (HE) and Elastica van Gieson histological stainings were performed according to standard protocols. For chromogenic immunohistochemistry, antigen retrieval was performed by rehydrating and boiling the slides in either Tris-EDTA buffer (pH9; 643901; Klinipath) for 10 min or in pronase (0,1% protease XIV, P5147-5G; Sigma) for 6 min at 37°C. Protein blocking was achieved using Normal Antibody Diluent (VWRKBD09-999; ImmunoLogic), followed by wash in PBS on a rocking table at 10 rpm. Primary antibodies, such as anti-CD31 (M0823, clone JC70A; DAKO) and anti-Collagen Type IV (C1926, clone COL-94; Sigma), were incubated for 1 hour at room temperature. Subsequently, primary antibody detection and chromogenic visualization was performed with BrightVision poly-HRP (DPVR110HRP; ImmunoLogic) for 30 min at room temperature and DAB (VWRKBS04-999; ImmunoLogic) or NovaRed (SK-4800; Vector) for 7 min at room temperature. After dehydration, slides were counterstained with hematoxylin for 1 min and enclosed with Quick-D mounting medium (7281; Klinipath). A selection of 5 representative original brightfield images per HE-stained section was loaded into the open source-imaging platform, Fiji (imageJ 64 Bit for Windows), and used for blinded histopathologic evaluation of cell nuclei density, using the specific cell counter plug-in. Relative vascularity was determined in 3 representative CD31-stained tissue sections, by CD31-based tissue segmentation using advanced Image Analysis software (inform 2.4.1; PerkinElmer).

### Fluorescent multiplex immunohistochemistry

Formalin-fixed, paraffin-embedded (FFPE) OMCs were sectioned following a perpendicular orientation with respect to the direction of the epidermal layer (Suppl. Fig. 3a). Tissue sections (6-µm thickness) cut at a distance of approximately 200µm were stained for fluorescent multiplex IHC. Immunofluorescent visualization was performed, as previously described^57^, with the Opal seven-color IHC Kit (NEL797B001KT; PerkinElmer) containing the fluorophores: DAPI, Opal 520, Opal 540, Opal 570, Opal 620, Opal 650 and Opal 690. Slides were boiled in Tris-EDTA buffer for antigen retrieval and removal of Ab-TSA complexes. Primary antibodies used are listed in Supplementary Table 2. A cocktail of monoclonal antibodies (moAbs, also called “Tumor marker”) directed towards two melanoma-associated antigens was added to each panel, including: anti-tyrosinase and anti-SOX10. After Ab staining, slides were counterstained with DAPI for 5 min and enclosed in Fluoromount-G (0100-01; Southern Biotech).

### Tissue imaging and quantitative digital analysis

Whole tissue slides were imaged using Vectra Intelligent Slide Analysis System (Version 3.03, PerkinElmer Inc.). This imaging technology combines imaging and spectroscopy to collect entire spectra at every location of the image plane. Images of single stained tissues for each reagent were used to build spectral libraries of the single dyes by using the Nuance Multispectral Imaging System (Version 3.0.2, PerkinElmer Inc.). These spectral libraries were used to unmix the original multispectral images obtained with the Vectra imaging system (Suppl. Fig 3c), to obtain an accurate and specific quantification of the cleaved Caspase 3 negative, CD45 positive (Cl. Cas3^-^ CD45^+^) signal. A selection of 10-20 representative original multispectral images was used to train the inForm Advanced Image Analysis Software (Version 2.4.1, PerkinElmer Inc.) for quantitative image analysis (tissue segmentation, cell segmentation and positivity score) as described previously^57,58^. All settings applied to the training images were saved within an algorithm allowing batch analysis of multiple original multispectral images of the same tissue. Vectra Review (Version 2.0.8, PerkinElmer Inc.) was used to select the areas for analysis; this consisted of the entire reconstructed tissue. Both background area and tissue area (Region Of Interest or “ROI”; Suppl. Fig 3b) were defined by fully-automated trainable tissue segmentation, based on morphological features and expression of DAPI (for efficient cell segmentation) and Tumor markers (for discrimination of areas containing tumour cells). A region of disinterest (ROD) was manually drawn in a qualitative manner over the stratum corneum, dermal border and injection site in order to reduce the auto-fluorescent signal caused by the structural characteristics of those regions. To appreciate the cell distribution patterns at low resolution, we clustered the cell positions using hierarchical mean linkage clustering, with a distance threshold of 150 mm (i.e., clusters whose centres were more than 150 mm apart were not joined). Single-cell-based information was saved to a file format compatible with flow and image cytometry data analysis software FlowJo (Version 10, Treestar).

### Time-lapse image acquisition and processing

All time-lapse experiments were performed using a multiparameter multiphoton microscope (TriMScope-II, LaVision BioTec, Bielefeld, Germany) on a temperature-controlled stage (37°C). 4D time-lapse recordings were acquired by sequential scanning with 950 nm (eGFP and PKH26) and 1090 nm (SHG). Areas of interest were imaged with 120sec time interval between individual scans over a 2-hour period, starting approximately 4 hours after pre-labelled cDC2 injection. Images were analyzed using the open source-imaging platform, Fiji (imageJ 64 Bit for Windows). Drifts in time-lapse recordings were corrected using the Correct 3D drift plugin. If necessary, images were scaled and adjusted for brightness and contrast to enhance visualization.

### Flow Cytometry and Flow Cytometry Cell Sorting (FACS)

A complete list of antibodies used in the study is reported in Supplementary Table 1. Dead cells were identified using Fixable Viability Dyes eFluor® 506 or 450 (Affymetrix, eBiosciences) and excluded from the analysis. For surface staining: cells were incubated in 2% human serum for the blocking of non-specific antibody binding to receptors (10 min, 4°C) and subsequently stained with directly labelled primary antibodies (30 min, 4°C) (Supplementary Table 1). For IL-6 intracellular staining: cells were stimulated with LPS (1μg/mL) for 6hrs and GolgiPlug (BD Biosciences) was added in the last 5hrs of the assay. After staining of surface markers (using directly labelled anti-CD45, anti-CD1c and anti-CD14 Abs), cells were washed, fixed and permeabilised with Cytofix/Cytoperm buffer (BD Biosciences) and stained with FITC-labelled anti-IL-6 (BioLegend). Acquisition was performed on a FACSVerse flow cytometer (BD Biosciences) and FlowJo analysis software (Treestar) was used for data analysis. Flow Cytometry Cell Sorting (FACS) was performed using the ARIA SORP (Becton Dickinson, Franklin Lakes, NJ). Anti-CD45, anti-CD1c and anti-CD14 sterile antibodies were used. Briefly, 2 days post injection tumour-free OSCs and OMCs were digested, under sterile conditions, and the cell suspensions filtered, stained and cells were sorted with >98% purity.

### Mixed lymphocyte reaction and activated T cell phenotype

The ability of cDC2s (CD1c^+^CD14^-^) and CD14^+^ DCs (CD1c^+^CD14^+^), isolated from OSCs and OMCs, to induce T cell proliferation was tested in a mixed lymphocyte reaction (MLR). A total of 3×10^4^ CFSE-labelled unstimulated allogenic CD3^+^ T cells from a healthy donor were seeded in a round bottom 96 well/plate and 15.000 (1:2 ratio), 6.000 (1:5 ratio) or 0/myeloid cells were added. The percentage of proliferated T cells was analyzed at day 5 after staining with anti-CD3 and anti-CD8 antibodies. For assessing the myeloid subsets stimulatory potential on autologous T cells, 1×10^5^ autologous unfractionated CD3^+^ T cells were cultured with 10.000 (1:10 ratio) or 0/myeloid cells. After 5 days T cell were stained with anti-CD3, CD8, CD25 and HLADR (Supplementary Table 1) and acquired at the FACSVerse (BD Biosciences).

### Gene expression analysis

Total RNA was extracted from FACS-sorted immune cells using Trizol (Invitrogen), and cDNA was generated using the SuperScript II Reverse Transcriptase Kit (ThermoFisher Scientific). cDNA was then used as a template for messenger RNA (mRNA) amplification using the TranscriptAid T7 High Yield Transcription Kit (Thermo Scientific). Amplified (aRNA) was purified using Agencourt RNAClean XP beads (Beckman Coulter, Brea, CA) prior to Nanodrop quantification. 500ng/mL of aRNA were used for reverse transcription with Superscript II (ThermoFisher Scientific). The resulting cDNA was used as a template. qPCR using FastStart SYBR Green Master (Roche Diagnostic GmbH) was performed on a CFX96 real-time cycler (Bio-Rad). The 2–ΔΔCt method, in which Ct represents the threshold cycle, was applied. Samples were run in triplicate. Relative gene expression was determined by normalizing the gene expression of each target gene to β-actin (ACTB). Primer sequences are listed in Supplementary Table 3.

### Statistical analysis

Statistical analysis was performed using GraphPad Prism V6. Values in the present study are expressed as mean ± s.d. or SEM as indicated in the figure legends. The significance between two groups was analysed by a two-tailed Student’s *t* test. For statistical analysis of mixed lymphocyte reaction (Fig. 5) comparing cDC2s (CD1c^+^CD14^-^) and CD14^+^ DCs (CD1c^+^CD14^+^), a one-way ANOVA plus Bonferroni multiple comparison post-test correction was performed. Statistical significance was annotated as follows: *p < 0.05, **p < 0.01, ***p < 0.001.

## Supplementary Figure Legends

**Supplementary Figure 1. Skin biopsy decellularization results in cell- and vessel-free de-epidermized human dermis, with a preserved structural matrix and basal layer.**

**a,** Representative image of IHC evaluation of cellular (hematoxilyn-eosin), endothelial (CD31) and extracellular matrix (Elastica van Gieson) components in pre- and post-decellularization skin samples. **b,** Representative stainings of basement membrane (BM, Collagen type IV) in native skin and de-epidermized human dermis. **c,d** Comparison of the quantitative assessment of dermal cell densities before and after decellularization. Nuclei (hematoxylin) were evaluated in 5 predefined areas (20x images) by independent histopathologic reviews using a digital imaging software. Dot plots of hematoxylin^+^ nuclei enumeration (**d**, “cellularity”); matched-colour symbols indicate different donors (n=3). **e,f** Extent of endothelial cells in pre-*versus* post-decellularization samples was determined as the percentage of CD31^+^ signal over the whole tissue area. Matched-colour dots in graph (**f**, “vascularity”) show replicate measurements of distinct donors (n=3).

**Supplementary Figure 2. Melanoma growth in human organotypic cultures can affect epidermal morphology.**

**a,** Representative hematoxylin-eosin staining showing how epidermal morphology is affected by high melanoma cell density. Different keratinocytes (KCs) and BLM melanoma cell ratios were co-seeded onto a de-epidermized human dermis to find the optimal KC:BLM ratio, which ensures complete KC growth and differentiation into a fully-developed epidermis. Determined optimal ratio was 25:1. Interspersed tumor nests are indicated with an asterisk. Scale bar 100µm.

**Supplementary Figure 3. Analysis of organotypic human skin melanoma culture sections by mean of fluorescent multispectral imaging**

**a,** Schematic representation of tissue sectioning for the analysis of immune cell distribution. **b,** Definition of the Region Of Interest (ROI) on the unmixed fluorescent IHC image containing DAPI (blue), CD45 (green), cleaved caspase 3 (Cl. Cas3, red), Tumor (yellow), used to perform qualitative and quantitative analysis in inForm. **c,** Spectral library from multispectral IHC representative image. Individual examples of Cl. Cas3 (Opal520), CD45 (Opal570), Tumor (Opal650), FAP (Opal690), and DAPI, as well as autofluorescence from an unstained section, were spectrally analysed to generate a spectral library in support of multispectral unmixing in a multiplexed assay.

**Supplementary Figure 4. Representative gating strategies for flow cytometry analysis of cellular phenotype**

**a,** Purity of freshly isolated cDC2s from peripheral blood of healthy donors. Highly pure CD1c^+^CD14^-^ (> 98%) cells were obtained from healthy donor PBMCs through magnetic cell sorting (MACS) using the CD1c (BDCA1) DC isolation kit combined with a pre-depletion step of monocytes using CD14-MACS microbeads. Cells were stained with primary directly-labelled antibodies: anti-CD1c, anti-CD14 and anti-CD20 Abs and the purity was assessed by flow cytometry. Representative dot plots of unstained and stained samples are shown. **b,** Representative gating strategy for the definition of single, live immune CD45^+^ and non immune CD45^-^ cells in digested OMCs.

**Supplementary Figure 5. CD14^+^DC percentage and phenotype in conventional co-cultures (2D) and in the organotypic melanoma culture**

Intra-donor comparison of cDC2s cultured with tumour cells (BLM, A375, Mel624) in conventional 2D co-cultures *versus* OMCs (n=7, over 4 experiments). **a,** Graph showing the percentages of cDC2s and CD14^+^DCs. Red asterisk indicate the only tumour condition (BLM) in which we detected a higher percentage of CD14^+^DCs compared to cDC2s. **b,** Histograms indicate ratio of percentages of CD14^+^DCs and cDC2s (P=0.0109). **c,** Heat map summarizing ratio of GeoMFI’s of CD14^+^DCs and cDC2s for the indicated markers (HLADR, CD163, PDL1, MerTK). Red: higher expression in CD14^+^DCs, blue: lower expression in CD14^+^DCs.

**Supplementary Figure 6. Functional characterization of CD14^+^DCs in OSCs and OMCs**

**a,** 2 days after total cDC2s injection, OSCs and OMCs were digested and CD1c^+^CD14^-^ (cDC2s) and CD1c^+^CD14^+^ (CD14^+^DCs) subsets FAC-sorted for RNA extraction and molecular characterization by qRT-PCR. Gene expression levels (2^-ΔCt^) for the indicated genes in cDC2s and CD14^+^DCs. ACTB was used as internal reference. **b,c** Proliferation of allogeneic (**b**) and activation of autologous (**c**) CD3^+^ T cells 5 days after co-culture with tumor-conditioned and tumor-free CD14^+^ DCs. T cells alone and cDC2s prior to tumour conditioning were used for comparison. **b**, Bar graphs (n of replicates = 3, means ± SEM) of two independent experiments are shown. **c,** T cells were stained for HLADR, CD25, CD3, CD8 and live/dead marker. Percentage of activated CD25^+^HLDR^+^ cells in total CD3^+^ T cells is reported. Bar graphs (n of replicates = 3, means ± SEM) of one representative experiment out of three is shown.

**Supplementary Figure 7. Gating strategy applied for the analysis of the patient tumor suspension.** Representative dot plots illustrating the gating strategy applied in the tumor suspension to identify: cDC2s (orange), CD14^+^ DCs (green), and CD14^+^ monocytes/macrophages (red) within total immune myeloid CD45^+^CD11c^+^ cells; cDC2s (orange), CD14^+^ DCs (green) in CD45+CD1c+ cells.

**Supplementary Figure 8. Culture of T lymphocytes in the OMC**

**a,** Intra-donor comparison of PBMCs cultured for 2 days in the OMCs *versus* conventional 2D culture tubes. For analysis, PBMCs were stained with anti-CD45, anti-CD3, anti-CD4 and anti-CD8 antibodies. After excluding doublets and dead cells, CD45^+^CD3^+^ cells were sub-gated in CD4^+^ and CD8^+^ cells.

**Supplementary Figure 9. Cell suspensions isolated from human melanoma lesions grown in a de-epidermized human dermis.**

De-epidermized human dermis was used as a scaffold for the growth of melanoma patient’s cell suspension. 2 days after injection, microtissues containing the patient material were fixed and processed for paraffin embedding. Multiplex IHC staining on tissue sections was performed using the following antibody panels: T cell panel (CD3, CD8, FoxP3, CD56, CD45RO and tumor marker), MDSC panel (CD68, CD163, CD15, HLADR, CD14 and tumour marker). Tumor marker consists of a mix of antibodies recognizing the melanoma antigens, HMB-45, Mart-1, Tyrosinase and SOX-10. Representative multicolor composite pictures for each panel are shown. Scale bar 50µm.

Author contribution
SDB and MT conceived the study, performed the experiments, analysed and interpreted the data, wrote the manuscript. GvW helped performing some of the experiments, analysing the data and writing parts of the manuscript. AvD and IS contributed to some experiments. MB arranged the collection of patient samples. MG and AV provided technical support. GB, SVH, JS and IJMdV contributed with critical feedback. EHvdB provided technical contribution and critical feedback. CGF conceived the study, interpreted the data and critically reviewed the manuscript.

## Aknowledgments

The authors gratefully acknowledge Rob Woestenenk and Gert-Jan Bakker for valuable assistance and technical support. This work was supported by Dutch Cancer Society grants KWO2009-4402 and 10673 and Netherlands Organization for Scientific Research Vici Grant 016.140.655. CF received the NWO Spinoza award and ERC Advanced Grant PATHFINDER (269019). MT was supported by an AIRC (Associazione Italiana Ricerca sul Cancro) Fellowship for abroad.

