## Supplementary Tables and Figures for "The tumour microenvironment shapes dendritic cell plasticity in a human organotypic melanoma culture"

**Supplementary Table 1. Flow cytometry antibodies**

| Marker | Clone | Isotype | Fluorochrome |
|---|---|---|---|
| CD1c | F10/21A3 | Mouse IgG1, κ | BV421 |
| CD14 | MϕP9 | Mouse IgG2b, κ | APCH7 |
| CD14 | MϕP9 | Mouse IgG2b, κ | PERCP |
| CD45 | HI30 | Mouse IgG1, κ | PERCP |
| CD45 | 5B1 | Mouse IgG1, κ | FITC |
| CD163 | GHI/6 | Mouse IgG1, κ | PE |
| MerTK | 590H11G1E3 | Mouse IgG1, κ | PECy7 |
| CD86 | 2331 (FUN-1) | Mouse IgG1, κ | PECy7 |
| HLADR | G46-6 | Mouse IgG2a, κ | FITC |
| HLADR | G46-6 | Mouse IgG2a, κ | PECy7 |
| CD11c | B-ly6 | Mouse IgG1, κ | FITC |
| CD11c | S-HCL-3 | Mouse IgG2b, κ | PE |
| CD11c | B-ly6 | Mouse IgG1, κ | APC |
| PDL1 | MIH1 | Mouse IgG1, κ | PECy7 |
| PDL1 | 29E.2A3 | Mouse IgG2b, κ | APC |
| CD206 | 15-2 | Mouse IgG1, κ | APC |
| CD206 | 19.2 | Mouse IgG1, κ | FITC |
| CD3 | SK7 | Mouse IgG1, κ | BV421 |
| CD4 | RPA-T4 | Mouse IgG1, κ | APCH7 |
| CD8 | SK1 | Mouse IgG1, κ | PECy7 |
| CD20 | L27 | Mouse IgG1, κ | FITC |
| CD25 | M-A251 | Mouse IgG1, κ | PE |
| IL-6 | MQ2-13A5 | Rat IgG1, κ | FITC |
| S100A9 | MRP 1H9 | Mouse IgG1, κ | FITC |

**Supplementary Table 2. IHC antibody details**

| **Marker** | **Clone** | **Isotype** | **Cat#** | **Supplier** | **Dilution** | **Antigen Retrival** |
|---|---|---|---|---|---|---|
| CD45 | 2B11+PD7/26 | Mouse IgG1, κ | M0701 | DAKO | 1:750 | EDTA 10’ |
| Cleaved Caspase 3 | 5A1E | Rabbit IgG | 9664 | Cell Signalling | 1/2000 | EDTA 10’ |
| FAP | SP325 | Rabbit IgG | M6250 | Spring Bioscience | 1/100 | EDTA 10’ |
| SOX10 | EP268 | Rabbit IgG | 383R | Cell Marque | 1/5000 | EDTA 10’ |
| HMB45 | HMB45 | Mouse IgG1 κ | M063401 | DAKO | 1/100 | EDTA 10’ |
| MelanA | A103 | Mouse IgG1 κ | MS-799 | Thermo Immunologic | 1/300 | EDTA 10’ |
| Tyrosinase | T311 | Mouse IgG2 a | MON10591 | Monosan | 1/200 | EDTA 10’ |
| CD3 | sp7 | Rabbit IgG | RM9107 | Thermo Scientific | 1/400 | EDTA 10’ |
| CD8 | C8/144B | Mouse IgG1 κ | M7103 | DAKO | 1/1600 | EDTA 10’ |
| CD45RO | UCHL-7 | Mouse IgG2 a | MS-112 | Thermo Scientific | 1/12460 | EDTA 10’ |
| Foxp3 | 236A/E7 | Mouse IgG1 κ | 14-4777 | eBioscience Affymetrix | 1/300 | EDTA 10’ |
| CD56 | MRQ-42 | Rabbit IgG | 156R-94 | Cell Marque | 1/500 | EDTA 10’ |
| CD163 | 10D6 | Mouse IgG1 κ | CD163-LCE | Leica | 1/500 | EDTA 10’ |
| CD68 | PG-M1 | IgG3 κ | M087601 | DAKO | 1/300 | EDTA 10’ |
| CD15 | MMA | IgM | 559045 | BD | 1/200 | EDTA 10’ |
| HLADR | LN3 | Mouse IgG2 b | MS-133-P1 | Thermo Scientific | 1/1260 | EDTA 10’ |
| CD14 | 7 | Mouse IgG2 a | NCL-L-CD14-2 | Novocastra | 1/60 | EDTA 10’ |

**Supplementary Table 3. Oligonucleotides Used for Quantitative RT-PCR**

F: Forward primer. R: reverse primer.

| Genes | Primer Sequence (5'-3') |
|---|---|
| *IL6* | F: GACAGCCACTCACCTCTTCAGAACG<br>R: ATCCATCTTTTTCAGCCATCTTTGG |
| *XBP1* | F: CCCTCCAGAACATCTCCCCAT<br>R: ACATGACTGGGTCCAAGTTGT |
| *THBS1* | F: TGCTATCACAACGGAGTTCAGT<br>R: GCAGGACACCTTTTTGCAGATG |
| *TLR8* | F: ggtcctctgctcagggtgtct<br>R: tgaatccagaaaacaaccacatg |
| *PTGS2* | F: gaatcattcaccaggcaaattg<br>R: ctgtactgcgggtggaacatt |
| *TLR4* | F: ggtgagtaattccatggtgcacta<br>R: ttccctctgcactggaagct |
| *SPP1* | F: gatagtgtggtttatggactgag<br>R: ttgtatgcaccattcaactcc |
| *IDO1* | F: TCTCATTTCGTGATGGAGACTGC<br>R: GTGTCCCGTTCTTGCATTTGC |
| *GZMB* | F: TGGGGGACCCAGAGATTAAAA<br>R: TTTCGTCCATAGGAGACAATGC |
| *HIF1A* | F: CACCACAGGACAGTACAGGAT<br>R: CGTGCTGAATAATACCACTCACA |
| *ACTB* | F: ctggaacggtgaaggtgaca<br>R: aagggacttcctgtaacaacgca |

SUPPLEMENTARY FIGURE 1

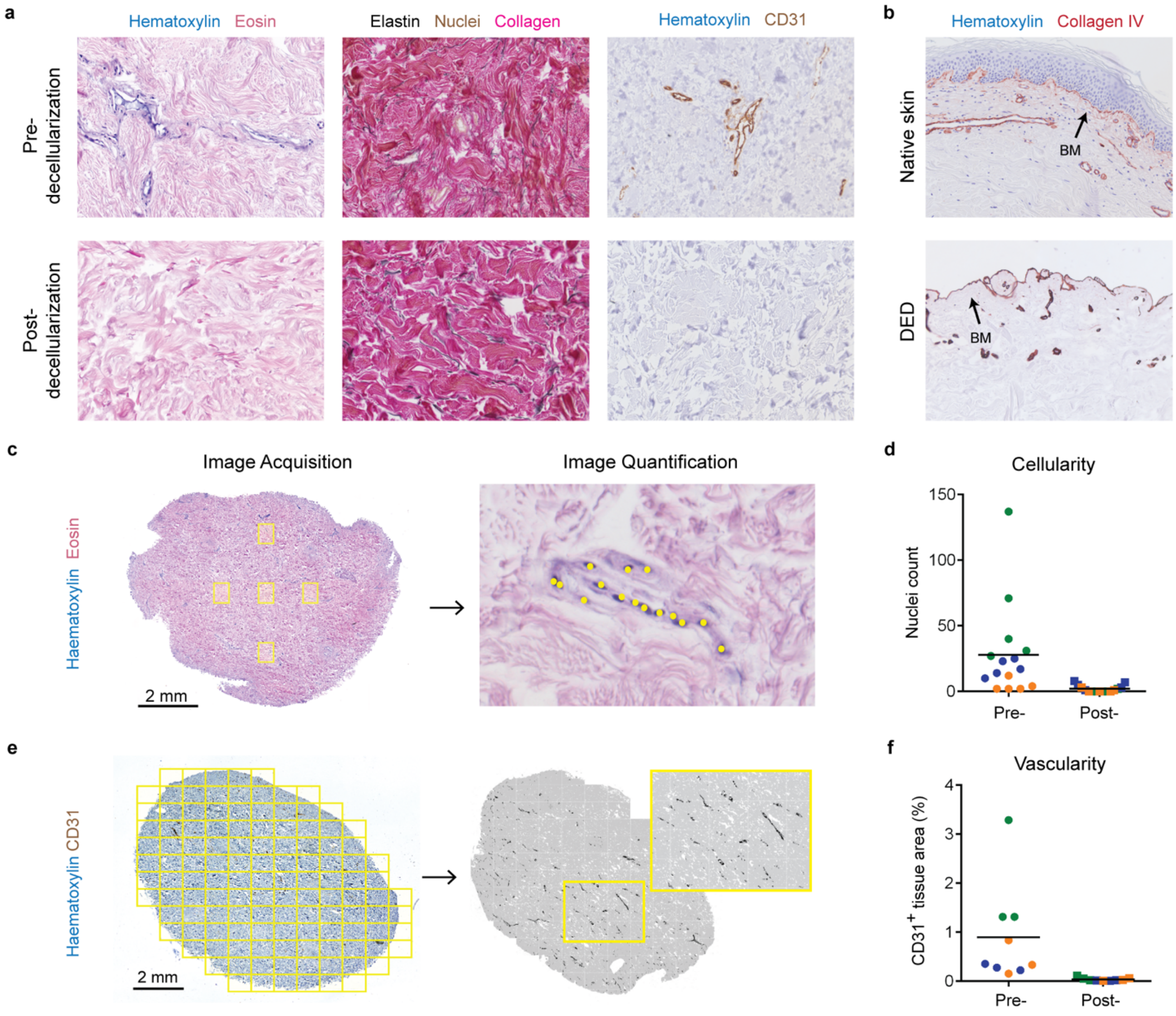

SUPPLEMENTARY FIGURE 2

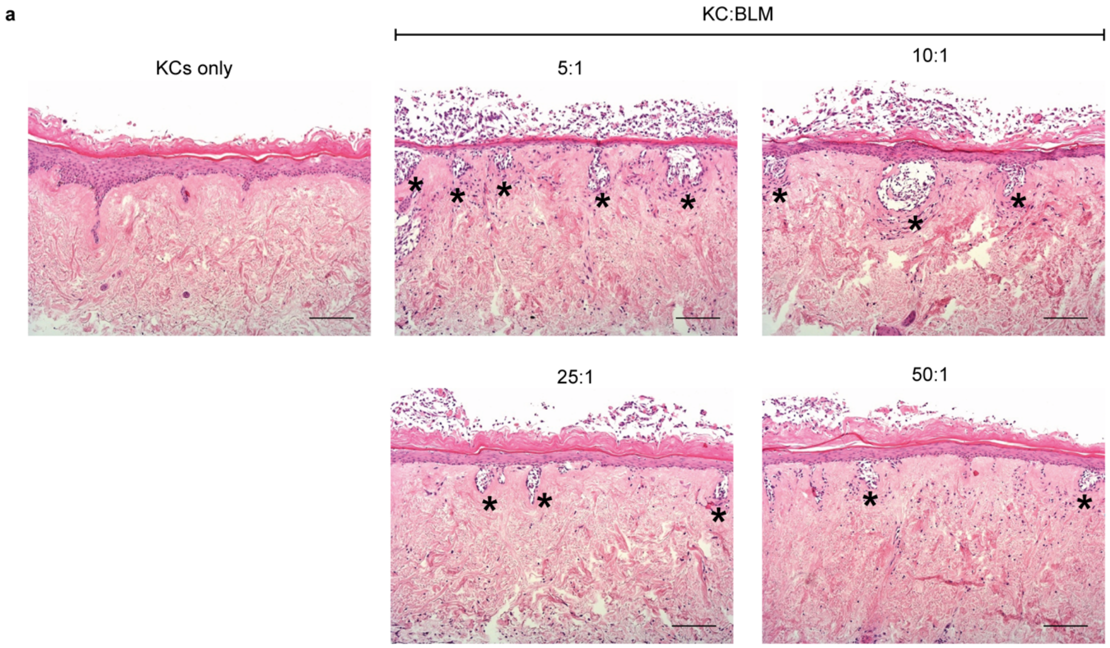

SUPPLEMENTARY FIGURE 3

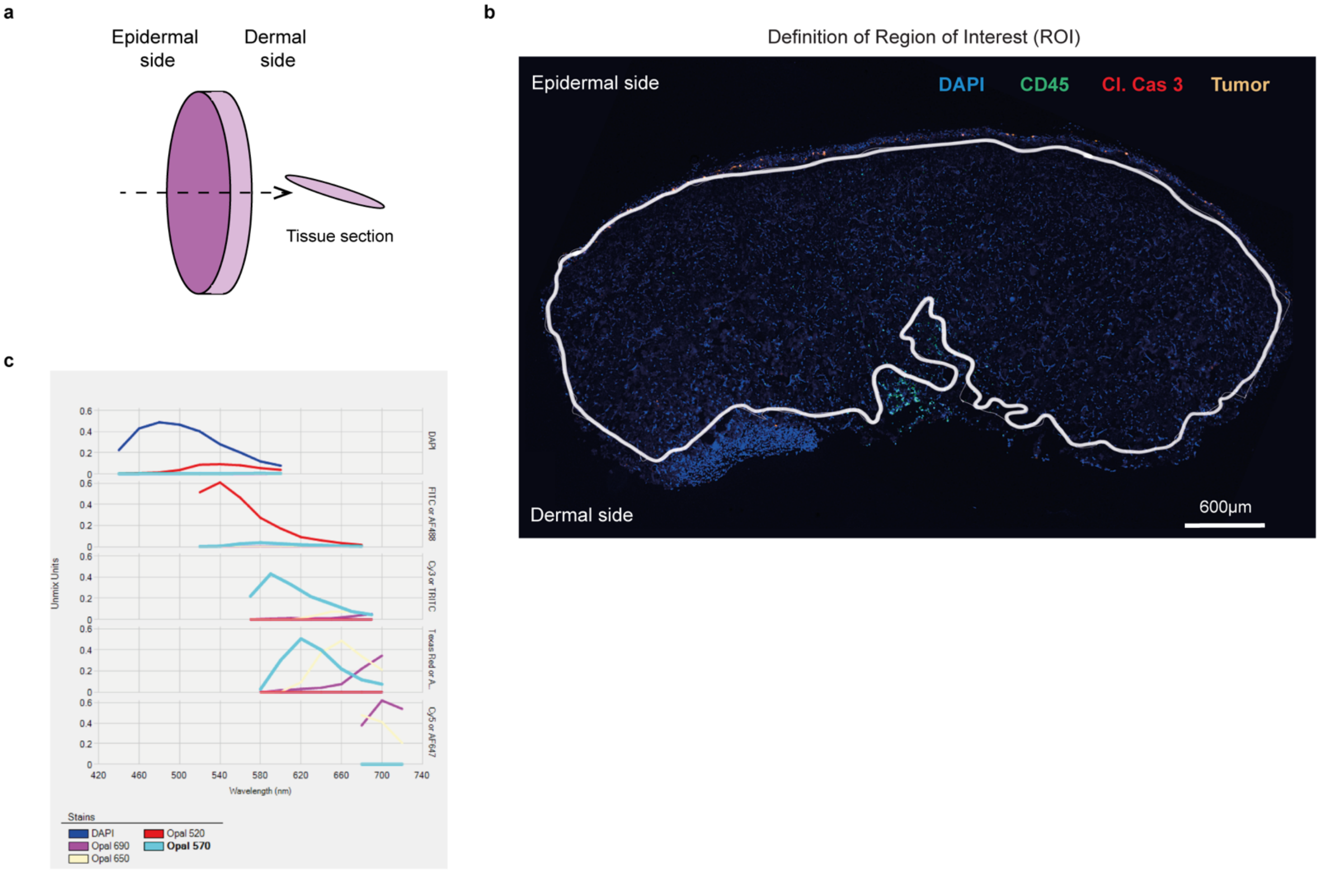

SUPPLEMENTARY FIGURE 4

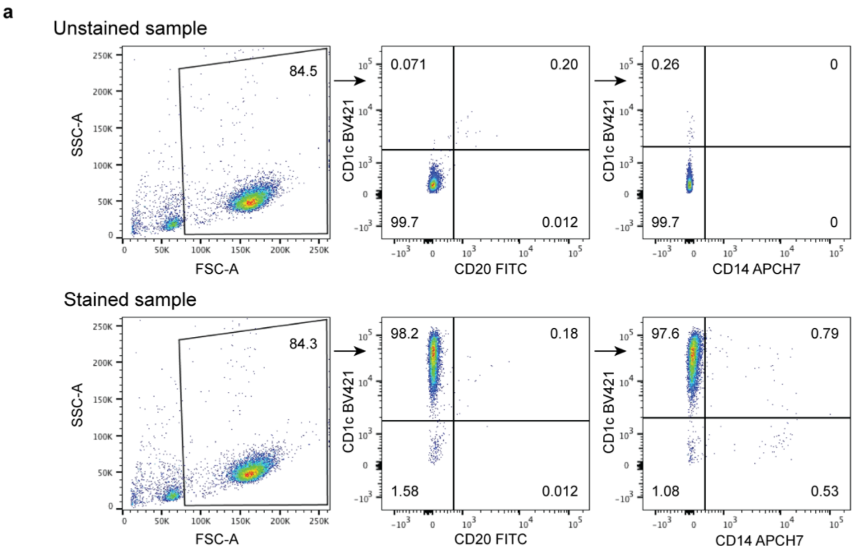


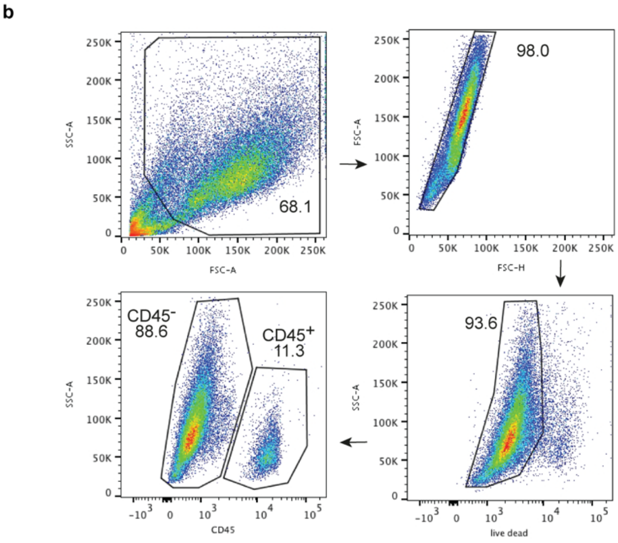

SUPPLEMENTARY FIGURE 5

**a**

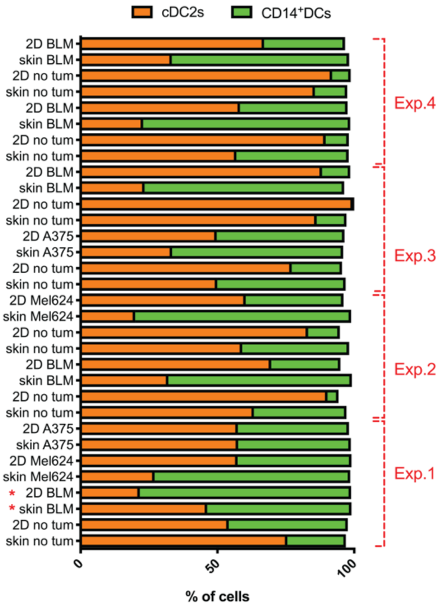


**b**

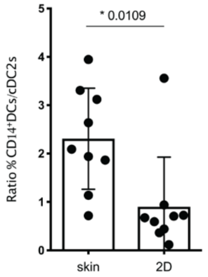


**c**

**Exp 1**

| | CD14⁺DCs/cDC2s BLM | | CD14⁺DCs/cDC2ss A375 | | CD14⁺DCs/cDC2s Mel624 | |
|---|---|---|---|---|---|---|
| | Skin | 2D | Skin | 2D | Skin | 2D |
| HLA DR | 0,2952 | 0,3770 | 0,2765 | 0,4273 | 0,4442 | 0,4238 |
| CD163 | 1,6289 | 1,4040 | 1,6667 | 1,1957 | 3,6165 | 1,2159 |
| PD-L1 | 1,9865 | 1,4603 | 1,6968 | 1,2828 | 1,7024 | 1,0776 |
| MerTK | 2,1831 | 0,7598 | 2,0262 | 1,3989 | 2,3315 | 1,3318 |

**Exp 2**

| | CD14⁺DCs/cDC2s BLM | | CD14⁺DCs/cDC2s Mel624 | |
|---|---|---|---|---|
| | Skin | 2D | Skin | 2D |
| HLA DR | 0,3545 | 0,1170 | 0,5064 | 0,3351 |
| CD163 | 2,2798 | 1,2999 | 1,9833 | 1,3665 |
| PD-L1 | 1,4491 | 0,7340 | 1,9431 | 1,3286 |
| MerTK | 1,6237 | 1,0664 | 1,9350 | 1,4759 |

**Exp 3**

| | CD14⁺DCs/cDC2s A375 | |
|---|---|---|
| | Skin | 2D |
| HLA DR | 0,6096 | 0,2636 |
| CD163 | 2,0365 | 1,2424 |
| PD-L1 | 0,9394 | 0,8145 |
| MerTK | 2,1339 | 1,2585 |

**Exp 4**

| | CD14⁺DCs/cDC2s BLM | |
|---|---|---|
| | Skin | 2D |
| HLA DR | 0,4270 | 0,2507 |
| CD163 | 1,6271 | 0,9843 |
| PD-L1 | 1,6636 | 0,6539 |
| MerTK | 0,7816 | 0,4727 |

SUPPLEMENTARY FIGURE 6

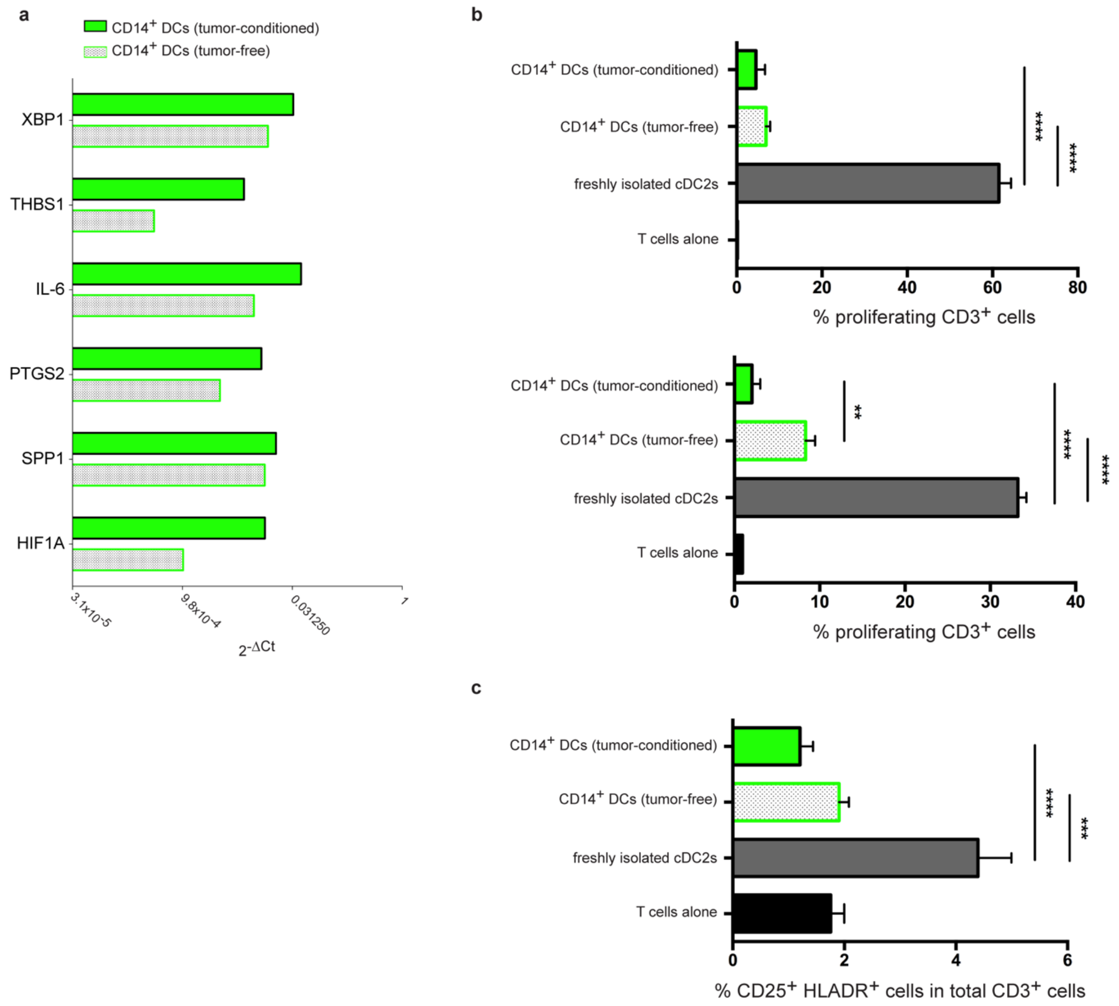

SUPPLEMENTARY FIGURE 7

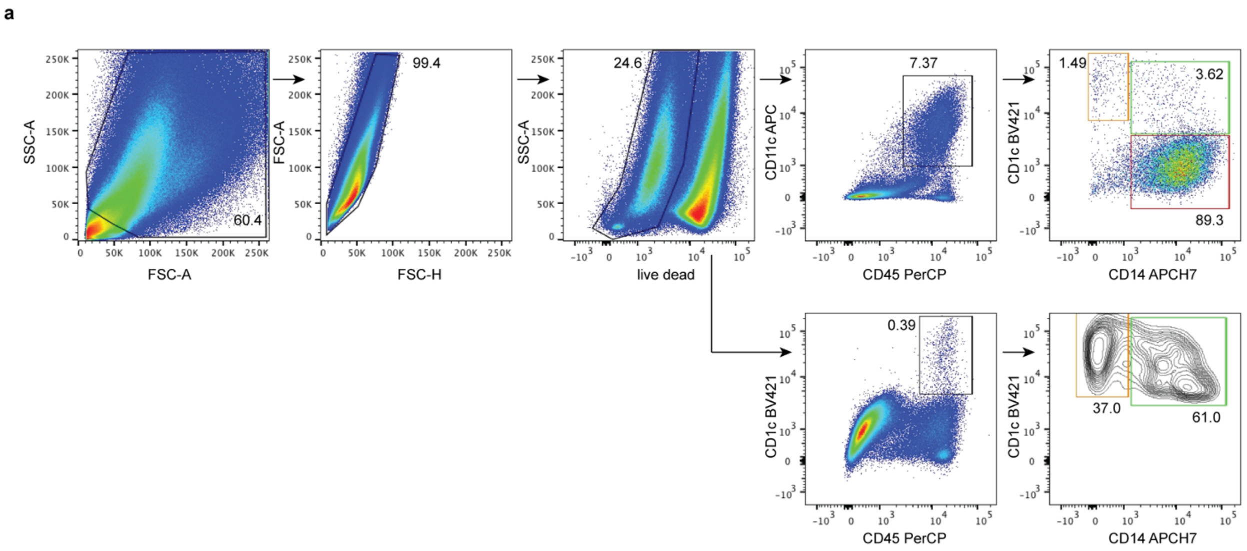

SUPPLEMENTARY FIGURE 8

**a**

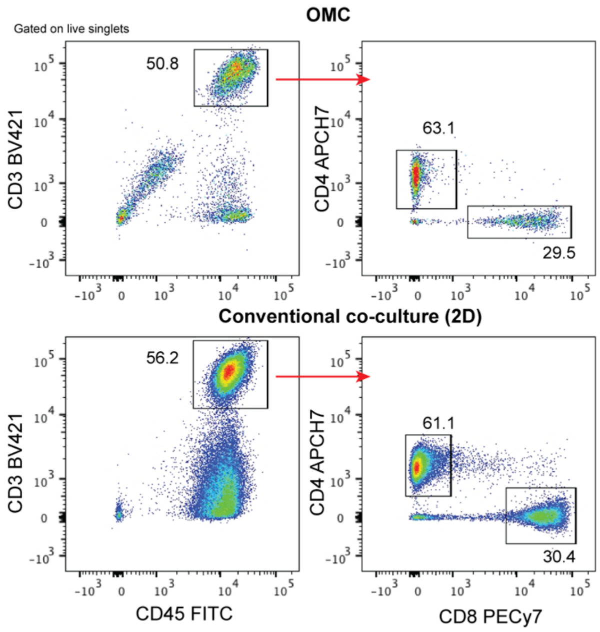


| Population | % of parental population | |
|---|---|---|
| | OMC | Conventional co-culture (2D) |
| $CD3^+$ | 50.8 | 56.2 |
| $CD4^+$ | 63.1 | 61.1 |
| $CD8^+$ | 29.5 | 30.4 |

SUPPLEMENTARY FIGURE 9

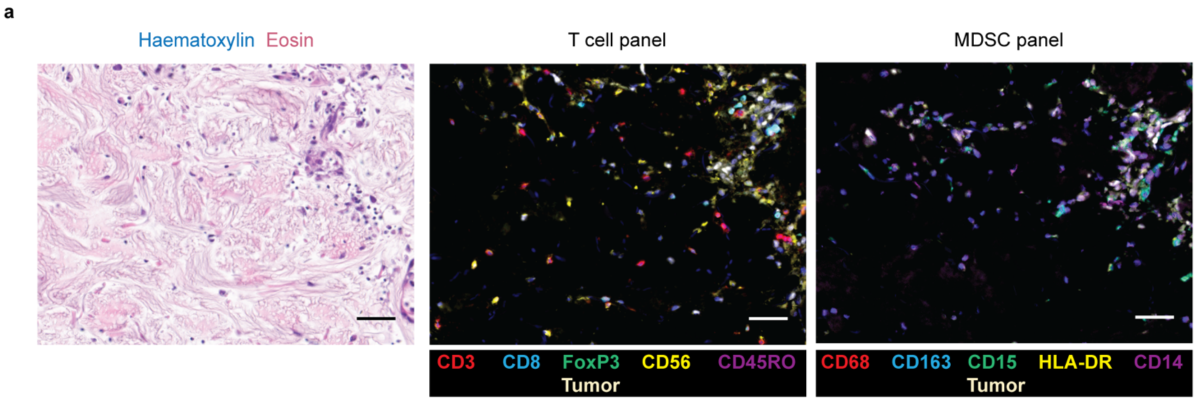
